# Decoding translational control by cis-regulatory elements and RNA binding proteins in human effector T cells

**DOI:** 10.64898/2026.05.05.722945

**Authors:** Branka Popović, Nikolina Šoštarić, Kaspar Bresser, Nandhini Kanagasabesan, Antonia Bradarić, Sander Engels, Aurélie Guislain, Mandy Rettel, Frank Stein, Joel I. Perez-Perri, Benoît P. Nicolet, Monika C. Wolkers

## Abstract

T cells are central players in killing virally infected and malignant cells. To achieve this, T cells depend on dynamic, tightly regulated alterations of the proteome. To decipher the rules instructing translation in T cells, we performed transcriptome and proteome analysis of polysome fractions from human effector CD8^+^ T cells. Transcriptome analysis informed on ribosomal occupancy of RNAs and uncovered the rapid and RNA-specific redistribution upon T cell activation. With machine learning, we identified RNA binding motifs that predict the RNA (re)distribution among the polysome fractions. Using matched proteome analysis, we identified polysome-associated RBPs and their swift shuttling across polysome fractions upon T cell activation. Integrating sequence feature analysis with polysome RBP localization uncovered PTBP1 as a positive translation regulator, through its interactions with cis-regulatory elements in 3′UTRs of target mRNAs. In conclusion, the multi-level analysis presented here identifies rules of selective translation control which shape the T cell proteome.

## Introduction

CD8^+^ T cells are key in our defense against pathogens and cancerous cells. Upon activation, T cells rapidly remodel their proteome, enabling them to differentiate, proliferate, and to produce cytotoxic molecules and pro-inflammatory cytokines, all of which is required for target cell elimination^1–3^. To enable such rapid responses, primed T cells maintain a state of readiness prior to activation^4–6^. At the same time, activated T cells require a tight regulation of their gene expression programs to prevent immunopathology^7,8^. Dysregulated gene expression is, amongst others, also associated with leukemias^9^. A better understanding of how T cells are wired is therefore paramount for developing more effective T cell-based immunotherapies for cancer, while offering therapeutic targets in autoimmune disorders.

T cell preparedness and activation-induced protein remodeling is intricately coordinated by transcriptional and post-transcriptional mechanisms^4,10,11^. Protein production depends on the availability of mRNA transcripts, whose cellular levels are shaped by transcriptional activity and post-transcriptional regulation, including splicing and mRNA stability. Intriguingly, the overall mRNA abundance is, with a correlation coefficient of ∼0.4 in mammalian cells, a poor predictor of the actual protein abundance^12–14^. Consequently, this finding indicates that the actual protein output from mRNA transcripts is highly regulated at the translational and post-translational level, as well as by protein turnover rates. Translation efficiency defines how many proteins are generated from a given mRNA molecule. Interestingly, T cell activation boosts ribosome biogenesis^6,15^, supporting the high demand on the translation machinery to massively increase protein synthesis upon T cell activation. Yet, protein production is highly selective and orchestrated and follows throughout time distinct programs^16,17^. How this selected protein production is achieved during early T cell activation, and which factors contribute to this process is to date, however, not well understood.

Post-transcriptional control is defined by cis-elements on the RNA, i.e. sequences on the transcripts that define the mRNA structure, and that contain motifs guiding the fate of mRNA ^18^. Cis-elements serve as binding hubs for trans-elements, i.e. micro-RNAs and RNA-binding proteins (RBPs)^18,19^. Dysregulation of RBP expression or function is implicated in many diseases^20,21^, demonstrating the importance of RBP-mediated post-transcriptional control. Evidence is also emerging around RBP-mediated translation control in T cells. For instance, ZFP36L2 sustains the preparedness of memory T cells by blocking translation of the key pro-inflammatory cytokines TNF and IFNG^5^. Other examples of selective RBP-mediated translation inhibition in T cells include that of ZFP36 and ATXN2L blocking translation of *IFNG* and *TNF*^2,22,23^, Roquin blocking *NFKBID*^24^, and PDCD4 blocking Glutaminase and SENP3^25^. RBPs can also promote translation, as exemplified by ELAVL1/HuR for cytokines during early T cell activation^2,26^, and by TIA1 and TIAL1, which promote the translation of TCF7, LEF1 and FOXP1 to sustain the naïve T cell state^27^. While the potential of a handful of RBPs in regulating translation in T cells is evidenced, a comprehensive study on RBPs regulating translation is to date lacking. Notably, T cells express over 1,500 RBPs^28–31^. Intriguingly, both the RNA binding profile of RBPs and their mode of action on the target mRNAs can substantially alter upon T cell activation^2,30^. Thus, to decipher the rules that govern RNA translation and the remodeling upon early T cell activation, it is paramount to systematically decipher the contribution of RBPs herein.

In this study, we measured the polysome-associated transcriptome and proteome in effector T cells. Using SONAR, a machine learning tool we developed to predict protein abundance based on sequence feature (SF) importance^13^, we identified SFs that inform on the RNA polysome profiles and the dynamic shifts in ribosome occupancy upon early T cell activation. Paired quantitative proteomics analysis revealed how RBPs with divergent annotations of function are distributed throughout the polysome fractions. By integrating the information of sequence feature importance with RBPome analysis, we identified RBPs regulating the ribosome occupancy of endogenous mRNAs. Specifically, whereas HNRNPC and SRSF1 increase their ribosome-association in activated T cells and are critical for T cell survival and effector function, PTBP1 associates with ribosomes to support the translation of essential cell cycle-promoting genes. Together, our study reveals key mechanisms underlying selective translational control during early T cell activation.

## Results

### Rapid transcriptome and proteome remodeling upon T cell activation

To capture the rapid changes of gene expression upon T cell activation, we first measured the transcriptome and proteome of total cell lysates from non-activated primary human CD8^+^ effector T cells, and from T cells that were reactivated for 2h with PMA-ionomycin (**Fig 1A**; *see Methods section*). RNA-sequencing analysis revealed that 10,89% of the total transcriptome was differentially expressed (DEG) within the first 2h, with 2892 (6,23%) transcripts significantly increased and 2185 (4,66%) transcripts decreased (**Fig 1B, Suppl Table 1**). Most DEGs (76%) were protein-coding (**Fig 1C**), and 18% were lncRNAs (**Fig 1C, Suppl Table 1**). As expected for this early time point of T cell activation^16^, the changes in the proteome were limited. Nevertheless, 141 (2,4%) of the 5840 detected proteins were differentially expressed, of which most (87) were upregulated (**Fig 1D, Suppl Table 2**).

**Figure 1:**
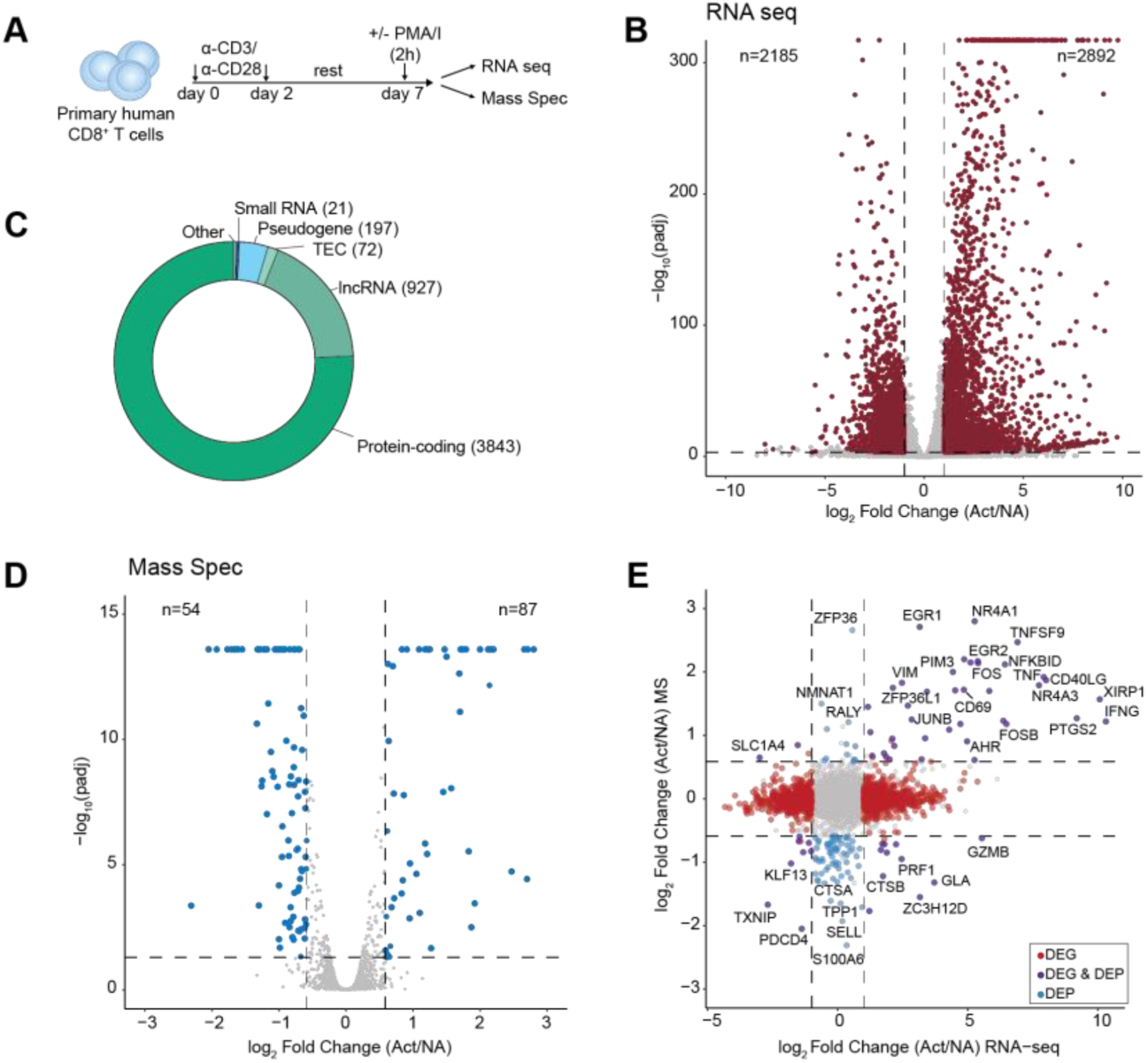
Effect of T cell activation on RNA/protein landscape. A) Experimental setup for transcriptome and proteome analysis from nonactivated and activated primary human CD8^+^ T cells. n=3 pools of 40 donors. B) Volcano plot showing differential RNA expression in nonactivated (NA) versus activated (Act) primary human CD8^+^ T cells, using the mean of 3 replicates. Differential expression was defined as log2 fold change (LFC)>1 and adjusted p-value < 0.001. C) Biotypes of the differentially expressed genes between nonactivated and activated T cells. D) Volcano plot showing differential protein expression in nonactivated (NA) versus activated (Act) primary human T cells, using the mean of 3 replicates. Differential expression was defined as LFC>1.5 and adjusted p-value < 0.05. E) LFC-LFC plot comparing RNA and protein differential expression analysis from nonactivated (NA) vs. activated (Act) T cells.

We next compared the transcriptome alterations upon T cell activation with proteome alterations. The log2 fold-change (LFC) of RNA abundance only mildly associated with changes in protein abundance (**Fig 1E**). Yet, altered protein abundance generally followed the changes in transcript levels (**Fig 1E**; *upper right and lower left quadrant*). This included the upregulation of effector molecules (TNF, IFNG), surface molecules (CD69, CD40LG, TNFSF9) and transcription factors (FOS, JUNB, AHR, NR4A1, NR4A3), and the downregulation of the translation repressor PDCD4, the transcription factor KLF13, and the thioredoxin-interacting protein TXNIP^6,15,32^ (**Fig 1E**). Interestingly, other proteins displayed increased (ZFP36, RALY) or decreased abundance (SELL/CD62L, TPP1, S100A6/Calcyclin) without apparent changes in mRNA abundance (**Fig 1E**, *upper and lower middle quadrant*), suggesting specific changes in mRNA translational output or protein turnover. Some proteins even showed opposing alterations in mRNA/protein abundance, which at least partially stemmed from protein secretion, as for Granzyme B (GZMB) and Perforin-1 (PRF1; **Fig 1E**, *lower right quadrant*), combined with increased *de novo* transcription upon T cell activation to replenish the protein pool at later time points^33^. In conclusion, T cell activation rapidly and substantially alters RNA and protein abundance in a controlled, targeted manner.

### Selective ribosome occupancy in effector T cells

To map early changes in translation in activated Teff cells, we performed polysome fractionation on matched samples from Fig 1 (*see Methods;* **Fig 2A**). We isolated RNA from 15 sucrose fractions, which based on 18S/28S ribosomal subunit distribution were pooled into ribosome-free (fractions 1-3), monosome-bound (fractions 4,5), polysome light (fractions 6-10) and polysome heavy-bound RNA (fractions 11-15; **Fig 2B**). Despite lower overall RNA content in polysome-heavy fractions compared to ribosome-free and monosome fractions (**,Suppl Fig 1A**), these fractions contained more distinct transcripts, with higher read counts (**Suppl Fig 1B**). To map the ribosome occupancy of each transcript, we compared abundance across polysome fractions. Because 1) the RNA composition of fractions varied (i.e., differing proportions of ribosomal RNA) and 2) transcripts within the polysome-heavy fraction were overrepresented in the RNA-sequencing libraries due to substantially lower total RNA content compared to other fractions (**Fig 2B**, **Suppl Fig 1A)**, standard comparative analysis based on normalized mRNA abundance was not possible. To overcome these limitations, we calculated the corrected mRNA abundance within each fraction (*see Methods*; **Suppl Fig 1C, Suppl Table 3**). These corrections allowed us to map the distribution of a given RNA across all fractions, without overestimating the RNA levels of the ribosome-denser fractions (**Suppl Fig 1D**).

**Figure 2.**
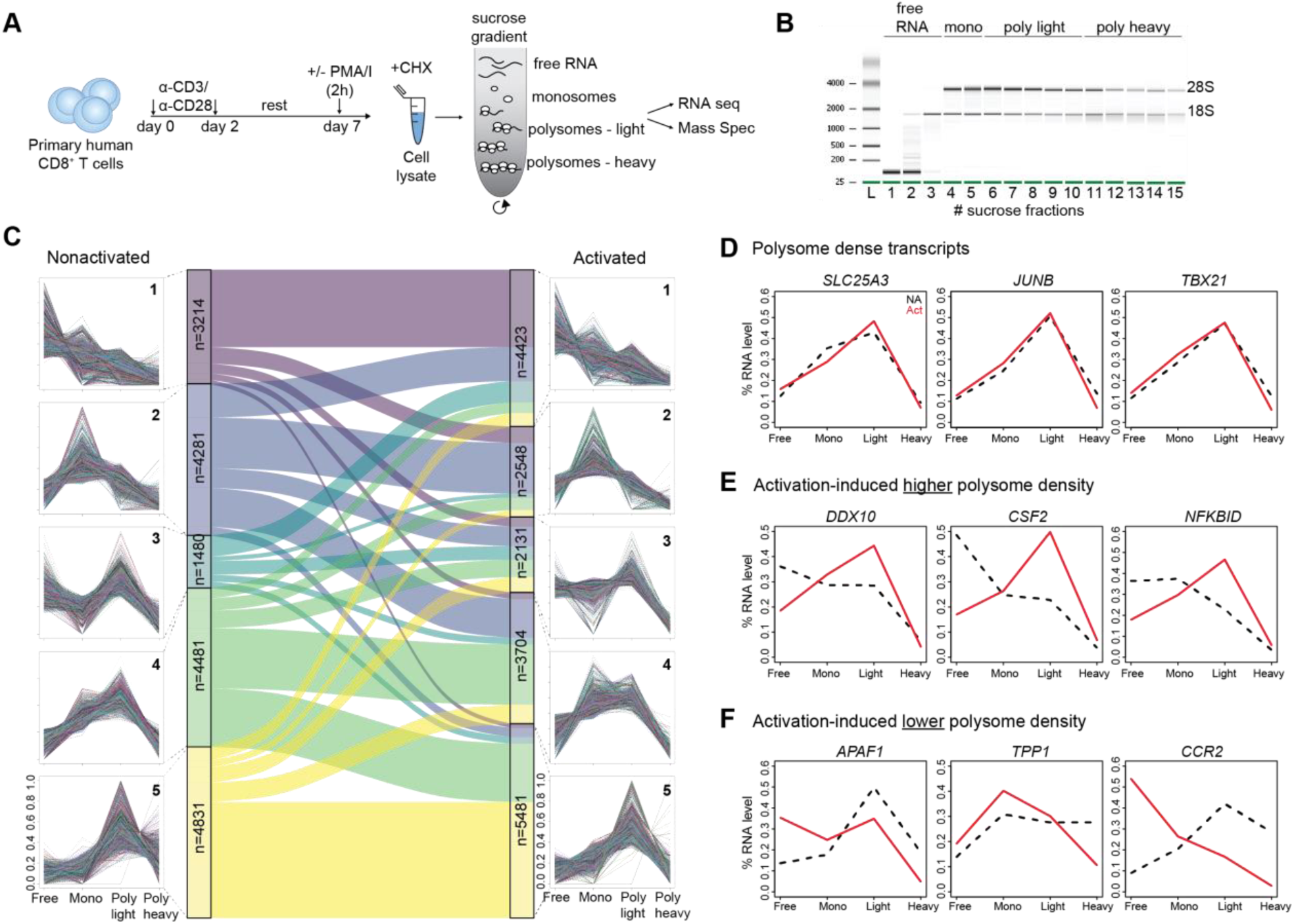
Selective association of RNA with polysomes in T cells depends on T cell activation. A) Experimental setup for polysome fractionation from nonactivated and activated primary human CD8^+^ T cells. n=3 pools of 40 donors. B) Total RNA distribution in collected sucrose fractions of activated T cells as determined by Bioanalyzer. C) Alluvial plot showing changes in RNA distribution profile across different clusters upon T cell activation. D-F) RNA distribution plots for indicated genes in nonactivated (NA) and activated (Act) CD8^+^ T cells.

To define changes in ribosomal occupancy upon T cell activation, transcript distribution curves throughout the polysome fractions were divided into 5 distinct clusters, from mostly ribosome-free (cluster 1) to mostly polysome-dense RNA (cluster 5; **Fig 2C**). Interestingly, even though many RNAs remain in the same cluster upon T cell activation, others substantial change their polysome profile (**Fig 2C, Suppl Table 4**). mRNAs with consistently dense polysome profiles included transcripts encoding the highly abundant solute carrier transporter SLC25A3^15^, TBX21 and JunB, two transcription factors critical for T cell effector function, the cell cycle proteins CCND2 and CDK6, and the DNA damage response protein CDK12 (**Fig 2D, Suppl Fig 1E**). Other transcripts increased their ribosomal occupancy upon T cell activation, including RNAs encoding the RNA binding protein DDX10, the cytokine GM-CSF (*CSF2*), and the NFKB Inhibitor Delta (*NFKBID*) (**Fig 2E**). mRNAs encoding the effector molecules IL2, IFNG and GZMB also slightly shifted towards higher ribosome occupancy upon T cell activation (**Suppl Fig 1F**). Conversely, transcripts encoding the apoptosome protein APAF1, the chemokine receptor CCR2, and the lysosomal protease TPP1 shown to reduce its protein expression upon T cell activation^6^ displayed reduced ribosome occupancy (**Fig 2F)**. Thus, early T cell activation results in dynamic and selective changes of ribosomal occupancy.

### Sequence features as determinants of RNA polysome profile in T cells

Because protein remodeling upon T cell activation is tightly controlled, we next aimed to decipher the rules that define ribosome occupancy. To this end, we employed SONAR, an XGBoost-based machine learning (ML) tool^13^. SONAR can predict >50% of the protein abundance in human immune cells based on sequence features (SFs) present in mRNA, such as UTR length, sequence conservation, % GC, putative RBP binding motifs and putative RNA modification sites (e.g. m6A, m7G)^13^. To employ SONAR models for translation regulation, we first calculated the translation efficiency for each RNA in T cells, as defined by the ratio of polysome-light and polysome-heavy abundance over its total RNA abundance (**Fig 3A**). Overall, translation efficiency spanned over a larger spectrum in nonactivated T cells, and it concentrated more towards the median in activated T cells (**Fig 3A**). To identify SFs that explain translation efficiency, we trained SONAR on three separate groups of RNA exerting different ribosomal occupancy (high: > 2; mid: 1.5 < x < 0.5; low: < 0.25) (**Fig 3A, B**). Models trained on random data had no predictive power (balanced accuracy of ∼0.33). In contrast, models trained to distinguish high, mid and low ribosome occupancy by SFs predicted >50% of the ribosome occupancy, irrespective of the T cell activation status (**Fig 3C**; balanced accuracy ranging from 0.53 to 0.67). To identify SFs contributing most to the models, we extracted the feature importance (*see methods*). In line with our previous findings^13^, feature importance depends on SF location within the mRNA, with the CDS region displaying the highest importance (**Fig 3D**). SFs located in the 5’UTR and 3’UTR contributed equally (**Fig 3D**). Interestingly, SF importance is higher in activated T cells than in nonactivated T cells, and this is observed in all mRNA regions (**Fig 3D**).

**Figure 3.**
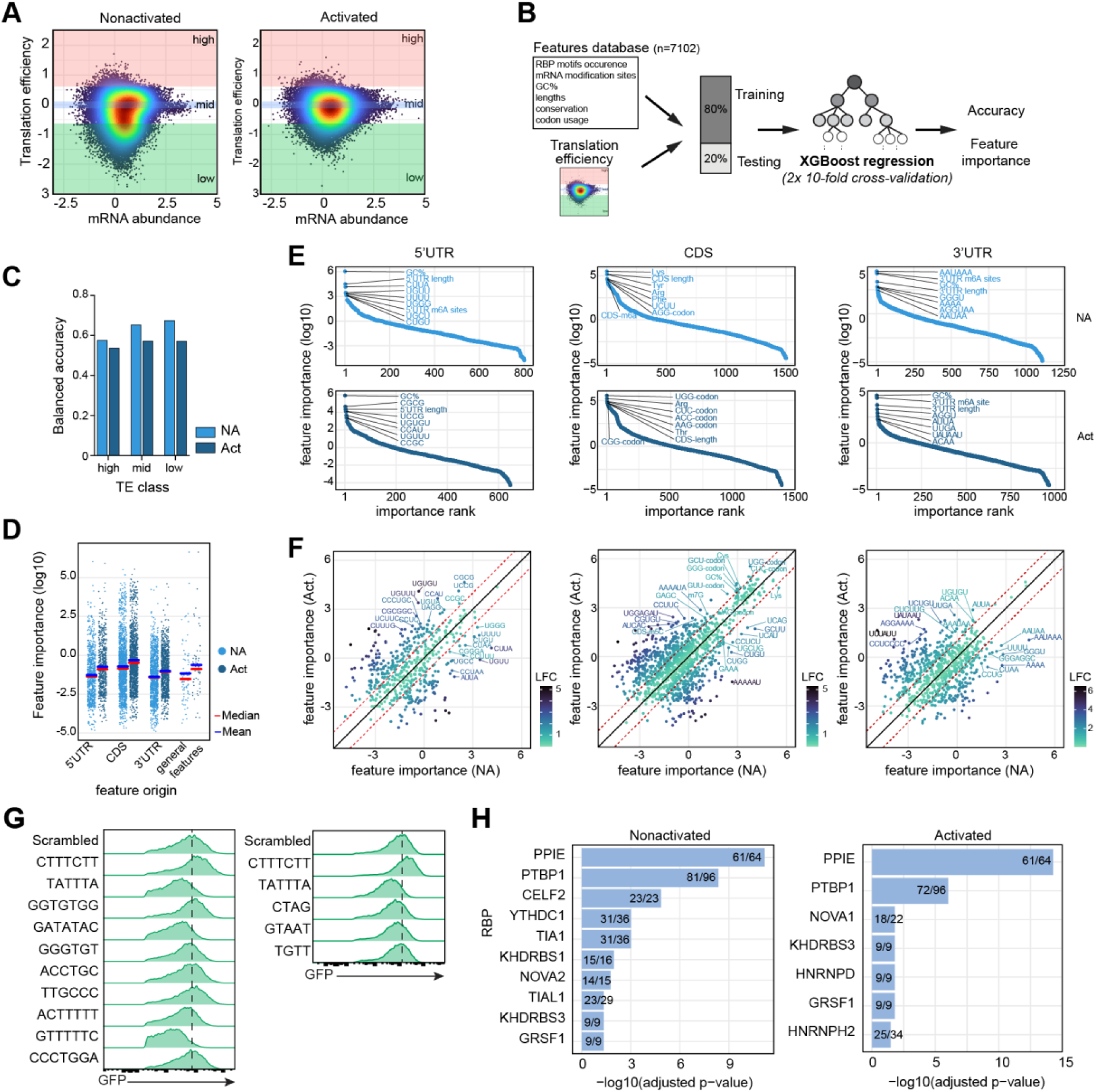
Sequence features predict RNA translation efficiency. A) Translation efficiency estimates (TE) were calculated by dividing the polysome heavy abundance by the total mRNA abundance (see methods) for nonactivated (left panel) and activated (right panel) T cells. Transcripts were split into 3 groups based on their translation efficiency estimates: low (TE <4), mid (0.75<TE<1.25), high (TE>4). B) Pipeline used to train and test SONAR models to classify transcripts into low, mid, high TE groups. C) Balanced accuracy of nonactivated (NA) and activated (Act) T cell SONAR models of (B). D) Feature importance of TE SONAR models in (B), separated by the feature origin (5’UTR, CDS, 3’UTR). Median (red) and mean (blue) are presented. E) Details of the top features in each origin (5’UTR, CDS, 3’UTR) for nonactivated (NA, top panels) and activated (Act, bottom panels). F) Comparison of feature importance of nonactivated (NA) and activated (Act) T cells, split per origins. G) Representative histograms of human CD8^+^ T cells transduced with GFP reporter constructs containing indicated motifs. H) Enrichment of RBP binding sites with importance >0 in 3’UTR in TE model. Fischer’s exact test. Graphs show RBPs with adjusted p-value < 0.05.

We next asked how individual SFs from the different mRNA regions contributed to the SONAR models. Irrespective of the T cell activation status, UTR length and the percentage of GC and GC-rich motifs contributed most to the 5’UTR models (**Fig 3E**, **Suppl Table 5, 6**). Individual GC- and CU-rich motifs, however, differentially contributed to models of nonactivated and activated T cells (**Fig 3E, F**). For the CDS, the region’s length, codon usage, and codon sets encoding for indicated amido acids belonged to the most important SFs (**Fig 3E**). Upon T cell activation, GC percentage, m7G modifications and the importance of codon sets encoding the amino acids Cysteine and Lysine altered for the CDS (**Fig 3E, F**). Top features for the 3’UTR included poly(A) signal (AAUAAA), putative m6A sites, GC content and 3’UTR length in nonactivated and in activated T cells (**Fig 3E**, **Suppl Table 5, 6**). Whereas AAUAAA (and derivatives) had a higher feature importance in nonactivated T cells, putative m6A sites, GC content, and 3’UTR length were equally important in nonactivated and activated T cells (**Fig 3E, F**).

Beyond translation efficiency, we questioned which SFs inform on the recruitment of ribosomes onto ribosome-free transcripts. To achieve this, we calculated the ribosome-bound to ribosome-free ratio (RF). On average, a lower RF was observed in activated T cells compared to nonactivated T cells (**Suppl. Fig2A**). In concordance with activation-induced protein remodeling^6,15,16,34^, the RF ratio in activated T cells was enriched for genes encoding proteins that regulate ribosome biogenesis, cytoplasmic translation and nucleosome organization (**Suppl Fig 2B**). SONAR models informing on SFs associated with the RF ratio reached an accuracy of 63% (**Suppl Table 7**). Intriguingly, SFs displaying high feature importance differed between DEGs and nonDEGs (**Suppl Fig 2C-D**). Furthermore, and concomitant with the previously reported potential of 3’UTRs to mediate post-transcriptional regulation^13,18,35^, SFs with high importance associated in particular with the 3’UTR (**Suppl Fig 2C-D**). In conclusion, mapping SFs associated with ribosome occupancy and translation efficiency identified targets that can be leveraged to manipulate protein expression in T cells.

### 3’UTR motifs with high feature importance in SONAR regulate protein production

Whereas SONAR models identify important SFs predicting ribosomal occupancy, they do not inform on the directionality of the translation regulation, i.e. high or low protein abundance^13^. We therefore experimentally tested 11 SF motifs with high feature importance identified in at least two SONAR models. We selected 3 short motifs (4-5 nt) and 8 long motifs (6-7 nt) (**Fig 3G**) and introduced six repeats of each motif in a 3’UTR-based GFP reporter system (*see Methods,* **Suppl Fig 2C)**. As controls, we included the CTTTCTT motif known to boost protein production^13^, and the TATTTA motif, an AU-rich element that effectively blocks protein production in T cells^5,14^. All 3 short motifs were suppressive, yet longer motifs displayed stronger, and more divergent effects on GFP protein expression (**Fig 3G**, **Suppl Fig 2D**). 3 out of 8 (GATATAC, ACTTTTT, GTTTTTC) strongly suppressed GFP protein expression (**Fig 3G**, **Suppl Fig 2D).** Intriguingly, the GTTTTTC motif suppressed protein expression even more effectively than the well-described ARE motif TATTTA (**Suppl Fig 2D**). In contrast, TTGCCC and GGTGTGG increased GFP expression, yet less so than the CTTTCTT motif (**Fig 3G, Suppl Fig 2D**). Possibly in part due to the long protein half-life of GFP, T cell activation did not reveal altered protein expression, except for the ARE motif TATTTA (**Suppl Fig 2E**). SONAR thus helps identify motifs within the 3’UTR that regulate protein expression in T cells.

### *In silico* analysis reveals highly prevalent putative RBP binding sites

Many SFs contributing to SONAR models can act as RNA-binding protein (RBP) hubs (**Fig 3B).** We therefore asked whether specific RBP motifs (as defined by the ATtRACT database^36^) were overrepresented among the predictive motifs. Enrichment analysis revealed high abundance of putative binding sites for PPIE, PTBP1, KHDRBS3 and GRSF1, independently of the T cell activation status (**Fig 3H**). Other RBP motifs were enriched in only one activation state, including CELF2, YTHDC1, TIA1 and TIAL1 in nonactivated T cells, and motifs for NOVA1, HNRNPD and HNRNPH2 in activated T cells (**Fig 3H).** Motifs for PTBP1, PPIE, CELF2 and TIA1 were enriched for both DEG and nonDEGs that displayed altered ribosome occupancy upon activation, and RBP hubs for RBMX was specifically enriched in DEG, and for YTHDC1 in nonDEG (**Suppl Fig 2H**). This *in silico* analysis identified RBPs that could potentially contribute to the ribosome occupancy and protein expression in T cells.

### Mapping polysome associated RBPs in resting and activated T cells

To experimentally measure the RBP distribution in polysome fractions, we performed mass spectrometry analysis on the identical samples we collected for RNAseq analysis (**Suppl Fig 3A, B**). Protein ranking analysis revealed that RBPs were highly abundant in each fraction (**Fig 4A**, *RBPs in red*). We identified in total 4834 proteins in the free RNA fraction, 4077 in the monosomal, 1568 in the polysome-light, and 847 proteins in the polysome-heavy fraction (**Fig 4B**). Interestingly, compared to 37% in the total cell lysate proteome, >60% of the proteins were RBPs in the polysome fractions (**Fig 4B**). Only a minority of the polysome-associated RBPs were ribosomal proteins, yet their abundance increased accordingly to ribosomal density (**Fig 4C**). The vast majority were other RBPs (**Fig 4C**). Examples include PDCD4 and ELAVL1/HuR that were detected in all fractions, ATXN2L in all fractions except polysome-heavy, and TIA1 found in free RNA and monosome fractions (**Suppl Table 8**). Of the RBPs with overrepresented putative binding sites *in silico (***Fig 3H**), we detected PTBP1 and YTHDC1 in polysome-light and PTBP1 also in polysome-heavy fractions, the latter independently of T cell activation (**Fig 4A, Suppl Table 8**).

**Figure 4.**
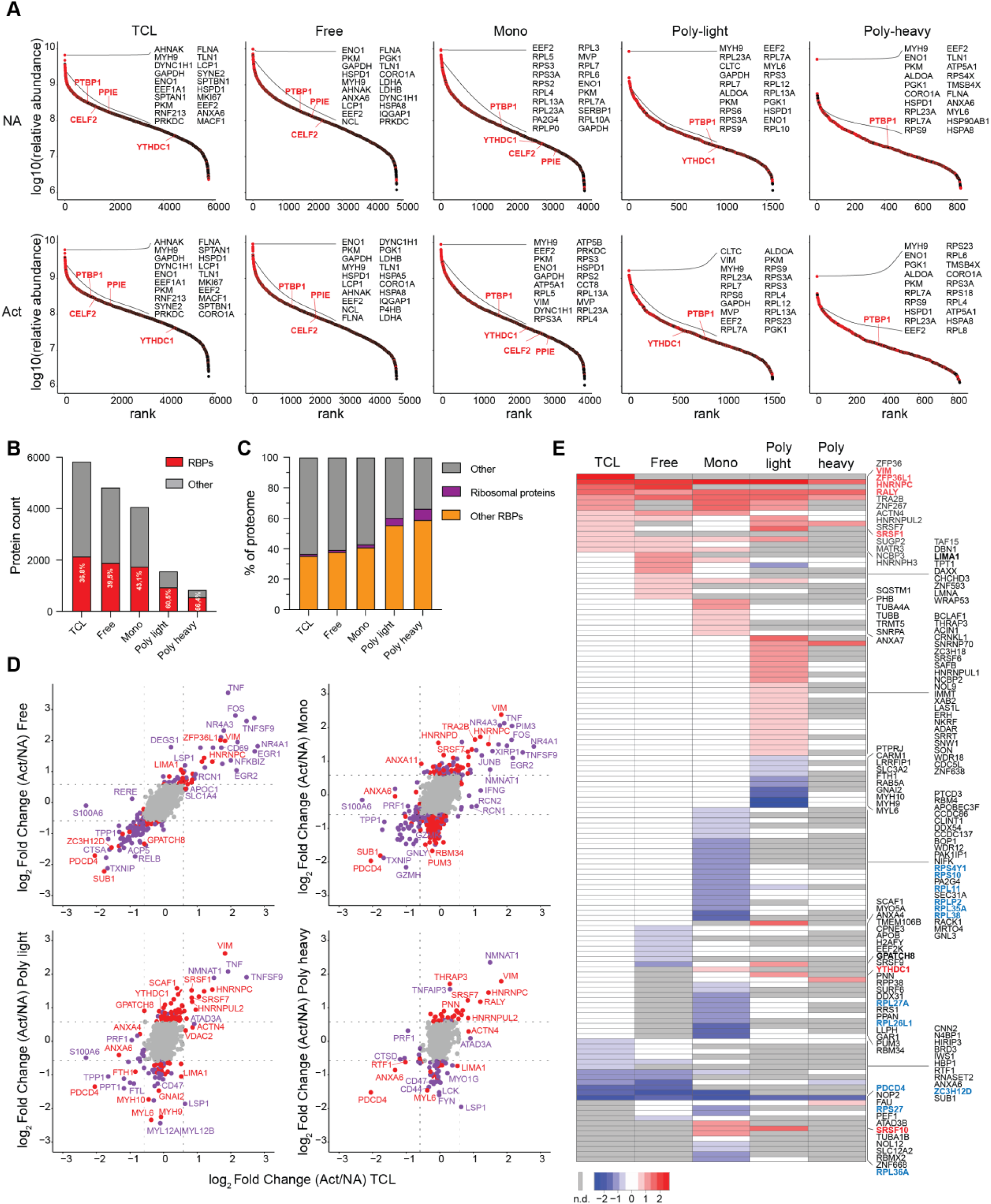
RBP enrichment to polysomes that redistributes upon T cell activation. A) Protein ranking and abundance of a proteins detected in total cell lysate (TCL) or indicated polysome fractions. Red dots represent RBPs. The top 20 RBPs and selected RBPs of the ones identified in Fig 3H are annotated. B) Number of identified proteins in MS analysis per indicated fraction. RBP count is indicated in red. C) Percentage of ribosomal proteins and other RBPs among all identified proteins of each fraction. D) Plots comparing LFC between proteins from nonactivated (NA) vs. activated (Act) T cells in total cell lysate proteome vs polysome fractions. RBPs with LFC>1.5 are indicated in red, and other proteins with LFC>1.5 in purple. E) Heatmaps showing differentially expressed RBPs in at least one fraction on the protein level. Centered log_2_ fold change (protein intensity Act/NA). Grey=not detected.

We next examined if polysome-associated RBPs altered their distribution upon T cell activation. The TMT6plex analysis (**Suppl Fig 3B**) allowed us to directly compare the protein content of each polysome fraction between nonactivated and activated T cells. Differential protein expression (DEP) analysis identified unique changes in protein expression for each polysome fraction (**Suppl Fig 3C; Suppl Table 8**). In line with the strong re-wiring of the cellular content, changes in the polysome-associated RBPs were substantial (**Suppl Fig 3C,** *in red*). Proteins upregulated in polysome-light and polysome-heavy fractions upon T cell activation were enriched for processes related to RNA metabolism and splicing processes (**Suppl Fig 3D**). Interestingly, the altered RBP expression in polysome fractions upon T cell activation did not seem to correlate with the overall RBP abundance in the total cell lysates. Rather, RBPs appeared specifically recruited to or expelled from polysome fractions, without changing its overall protein abundance (**Fig 4D**). A clear exception to this rule was PDCD4, which is not only released from ribosome-bound fractions, but is also actively degraded (**Fig 4D**), as reported^37^.

To further analyze which RBPs alter their association with polysomes and thus possibly contribute to translation control during early T cell activation, we mapped their significant increases or decreases upon activation (p-adjusted <0.05, LFC >1.5; **Fig 4E**). A subset of RBPs displaying up-or downregulated abundance in the total cell lysates were overall also more (VIM, HNRNPC, and RALY) or less (PDCD4) abundant in polysome fractions (**Fig 4E**). The majority of DE-RBPs, however, was up- or down-regulated in selected polysome fractions upon T cell activation, without displaying overt differences in protein expression in the total cell lysate (**Fig. 4E**). For instance, increased protein expression of SRSF1 upon T cell activation specifically resulted in its accumulation in the poly-light fraction (**Fig 4E**). Furthermore, even though the protein expression of ZFP36L1 rapidly increased upon T cell activation^2^ (**Fig. 4E**), it specifically accumulated in the ribosome-free fraction, with undetectable levels in polysome-containing fractions (**Fig 4E**). Similarly, downregulated protein expression of ZC3H12D resulted in specific reduction of its presence in the ribosome-free fraction (**Fig 4D, E**). Most RBPs, however, shifted between fractions upon T cell activation without changing their overall protein abundance. Interestingly, this included the ribosomal proteins RPS4Y1, RPS10, RPL11, RPLP2, RPL35A, RPL38, RPL27A, RPL26L1, RPS27 and RPL36A, displaying a specific reduction in the monosome fraction (**Fig 4E**). Other proteins like actin-binding LIMA1 moved from polysome-light to polysome-free fraction, and the splicing factor GPATCH8^38^ moved in the opposite direction (**Fig 4D, E**). YTHDC1 was also increased in the poly-light fraction (**Fig 4E**). Combined, this analysis demonstrates that RBP interactions with polysome fractions are highly dynamic.

### PTBP1, HNRNPC, and SRSF1 modulate translation and T cell fitness

We next asked whether and how polysome-associated RBPs modulate the protein expression in T cells. We focused on three proteins: PTBP1, which we identified as a particularly strong candidate regulator of RNA-polysome association (**Fig 3H, Suppl Fig 2H, Fig4A)** and HNRNPC and SRSF1 which displayed increased protein abundance and polysome association upon T cell activation (**Fig 4D, E**). To test their involvement in global translation, we measured puromycin incorporation^39^ upon CRISPR/Cas9 gene editing of these three proteins in primary T cells (**Suppl Fig 4A)**. Whereas the puromycin incorporation in HNRNPC KO and SRSF1 KO T cells was inconsistent (**Fig 5A, B**), PTBP1 KO T cells showed significantly reduced puromycin incorporation levels compared to control T cells, irrespectively of the T cell activation status (**Fig 5A, B)**. Thus, PTBP1 supports global translation in T cells.

**Figure 5.**
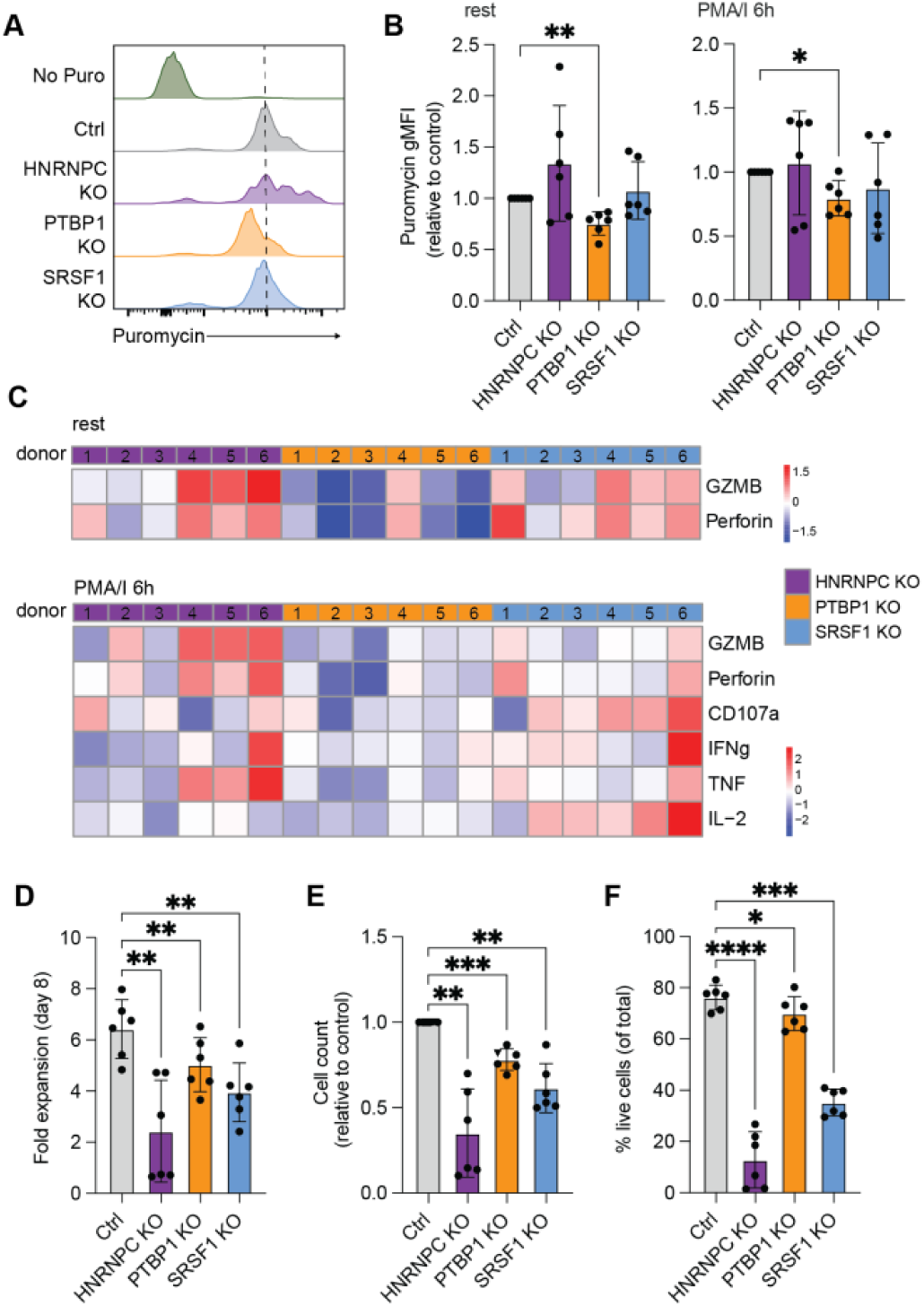
Polysome-associated RBPs differentially regulate T cell function. A) Representative histograms and B) Fold change of Puromycin gMFI levels in human RBP KO CD8^+^ T cells compared to controls. Data are presented as mean ± SD of 6 donors, compiled from 2 independently performed experiments. Repeated measures one-way ANOVA with Dunnett’s multiple comparison (*P<0.05, **P<0.01). C) Heatmaps of log2FC of gMFI of the indicated protein in RBP KO CD8^+^ T cells compared to controls. Numbers indicate the corresponding donor. D) Fold expansion and E) Total cell count of RBP KO CD8^+^ T cells compared to control treated T cells at 8 days after nucleofection with Cas9-RNPs. Compiled data of 6 donors from 2 independently performed experiments. Data are presented as mean ± SD. Repeated measures one-way ANOVA with Dunnett’s multiple comparison to control (**P<0.01, ***P<0.001). F) Viability of RBP KO T cells presented as percentage of live from total cells. Data are presented as mean ± SD. Repeated measures one-way ANOVA with Dunnett’s multiple comparison to the control (*P<0.05, ***P<0.001, ****P<0.0001).

We next defined the effect of RBP deletion on the T cell activation and effector function. Only HNRNPC KO T cells showed significantly reduced CD25 expression, yet increased CD69 and CD137 expression (**Suppl Fig 4B-D**). The cytokine production (IL2, TNF, IFNG) and expression of cytotoxic molecules (GZMB, Perforin), and degranulation marker CD107a was variable between donors (**Fig 5C, Suppl Fig 4E-N**). Yet, whereas HNRNPC KO T cells consistently expressed more Granzyme B, Perforin and CD107a (**Fig 5C, Suppl Fig 4M, N**), SRSF1 deficiency consistently increased IL-2 production, both in terms of percentages of IL-2^+^ T cells and IL-2 production per cell (**Fig 5C, Suppl. Fig4E, G**). As previously reported^40^, PTBP1 deficiency resulted in reduced Granzyme B expression (**Fig 5C, Suppl Fig 4K**). Importantly, even though all three RBP KOs affected T cell expansion and cell count, cell viability was in particular affected in HNRNPC KO and SRSF1 KO T cells (**Fig 5D, E**). Because low cell viability rendered data interpretation in follow-up experiments more challenging, we continued our studies with PTBP1, which displayed strong effects on the translation rates, and for which we identified many putative binding sites for polysome-associated mRNAs.

### A massively parallel reporter assay reveals PTBP1-mediated regulation of protein expression

To systematically examine how PTBP1 regulates protein expression through 3’UTR motifs, we employed a massively parallel reporter assay (MPRA) with a library of 467 synthetic 3’UTRs fused to GFP, each containing six occurrences of a 6-7 nucleotide oligomer^13^. After retroviral transduction of the MPRA library into activated CD8^+^ T cells, we deleted PTBP1 or used non-targeting RNPs as control (**Fig 6A**). After 6 days of rest, we reactivated PTBP1 KO and control T cells for 16h with α-CD3/α-CD28, or left them nonactivated. The 15% top (GFP^hi^) and 15% bottom (GFP^lo^) GFP-expressing cells were FACS-sorted, and motif enrichment for high or low protein expression was determined by sequencing (**Suppl Fig 5A**). 13 putative PTBP1 binding sites present in the library were generally enriched in the GFP^hi^ population of control-treated T cells compared to “non PTBP1 motifs” (**Fig 6B**, *grey*). Importantly, this enrichment of PTBP1 motifs in GFP^hi^ T cells was partially lost in the absence of PTBP1 (**Fig 6B**, *orange*), highlighting that the MPRA identified PTBP1-dependent regulatory sequences. This included the canonical PTBP1 motifs CTCTCT and GTCTTT, independently of the T cell activation state (**Fig 6C**). Intriguingly, several motifs were sensitive to PTBP1 deletion only in the context of a specific T cell activation state. The effect on GFP expression of the CTTTCTT motif, which we previously identified as one of the strong positive regulators of protein expression^13^, was abolished in PTBP1 KO only upon T cell activation (**Fig 6C**). Conversely, the shift of the ATCTTC motif towards the GFP^low^ population was only observed in nonactivated T cells (**Fig 6C**). We also found additional putative PTBP1 motifs not yet annotated in the ATtRACT database, such as AGATAT, which repressed upon PTBP1 deletion, and ATTAAT which lost its repressive capacity in PTBP1 KOs (**Suppl Fig 5B**). Combined, MPRA confirmed known PTBP1-binding motifs and defined their effects on protein expression in primary T cells. In addition, we identified potential novel PTBP1 binding motifs and characterized their activation-dependent regulation.

**Figure 6.**
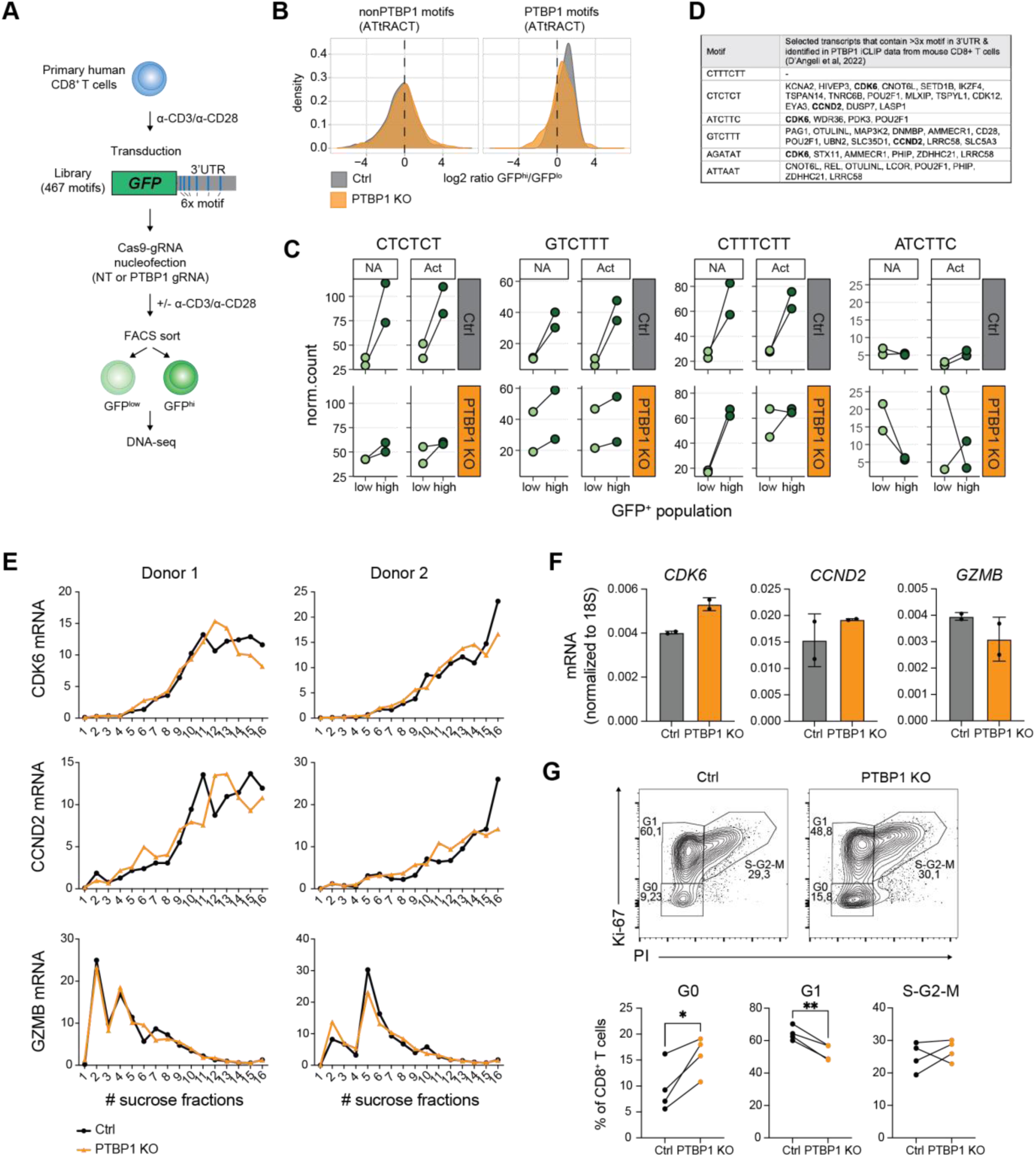
Motif-specific regulation of translation by PTBP1. A) Experimental setup for MPRA screen with GFP-3’UTR library containing 467 motifs in PTBP1 KO and control CD8^+^ T cells (NT = non-targeting). n=2 pools of 5 donors. B) Histograms depicting Log_2_ ratio of known PTBP1 motifs (right histogram) or other motifs (left histogram) between GFP^hi^ and GFP^low^ CD8^+^ T cells in PTBP1 KO and control. C) Graphs depicting normalized counts of indicated motifs in GFP^hi^ and GFP^low^ sorted CD8^+^ T cells. D) Table showing transcripts that are enriched for at least 3 repeats of indicated PTBP1 motifs in their 3’UTR and that were reported PTBP1 targets in mouse CD8^+^ T cells. E) Polysome fractionation of PTBP1 KO and control CD8^+^ T cells. Graphs show *CDK6*, *CCND2* and *GZMB* mRNA distribution through all 16 collected fractions. F) *GZMB*, *CDK6* and *CCND2* mRNA levels in PTBP1 KO and control CD8^+^ T cells. Data are presented as mean ± SD of 2 donors. G) Representative plots and quantification of cell cycle phase based on Ki-67 and PI staining in PTBP1 KO CD8^+^ T cells and control at 48h after activation with α-CD3/α-CD28 (n=4 donors). Paired t-test (*P<0.05, **P<0.01).

### PTBP1 regulates ribosome occupancy of endogenous mRNAs

Finally, we tested whether PTBP1 deficiency modulated polysome occupancy of endogenous mRNAs. We focused on mRNAs containing >3 repeats of a given PTBP1 motif in their 3’UTR, and that were experimentally confirmed PTBP1 targets in mouse CD8^+^ T cells^40^ (**Fig 6D**). Amongst the transcripts meeting these requirements were two genes involved in cell cycle, CDK6 and CCND2 (**Fig. 6D**). To determine whether PTBP1 regulates their ribosome occupancy, we performed polysome fractionation of PTBP1 KO and control CD8^+^ T cells. Both *CDK6* and *CCND2* mRNA are polysome-dense transcripts (**Fig 6E**). In the absence of PTBP1, however, both mRNAs displayed reduced ribosome occupancy (**Fig 6E**). In contrast, *GZMB* mRNA, another PTBP1 target^40^ that showed reduced protein expression in PTBP1 deficient T cells (**Suppl Fig 4K**), but that does not contain these regulatory elements in its 3’UTR, displayed limited polysome binding, and its polysome profile did not alter in PTBP1 KO T cells (**Fig. 6E**). Interestingly, when measuring the total mRNA levels of the PTBP1 target genes, *GZMB* mRNAs appeared slightly lower, and *CDK6* mRNA levels were higher in PTBP1 deficient T cells (**Fig 6F**), suggesting additional, target gene-specific effects of PTBP1. Importantly, consistent with the reported effects of CDK6 and CCND2 on the cell cycle^41,42^, the effect of PTBP1 on CDK6 and CCND2 translation corroborated with a higher percentage of PTBP1 KO T cells in the G0 phase and a lower percentage in the G1 phase (**Fig 6G**). We therefore conclude that PTBP1 regulates the ribosome occupancy of endogenous target mRNAs in effector CD8^+^ T cells in a sequence-specific manner.

## Discussion

In this study, we provide the first comprehensive analysis of translation regulation in early human CD8^+^ T cell activation. By integrating polysome-associated transcriptome and proteome profiling, we uncover cis- and trans-acting regulatory features that define the dynamic, activation-dependent association of RNAs and proteins with ribosomes. Combined, our study provides critical insights in the contribution of RBPs coordinating translation regulation in T cells.

We identified PTBP1 as a positive regulator of translation in T cells. PTBP1 binding sites were predicted by SONAR, and their role in defining protein abundance through PTBP1 was confirmed by an MPRA assay. In line with the reduced cell expansion in PTBP1 KO T cells, endogenous *CDK6* and *CCND2* mRNA containing PTBP1 binding motifs had less dense polysome profiles in the absence of PTBP1. Thus, in addition to its reported role in T cell activation and B cell development by regulating alternative splicing and RNA stability^40,43–45^, we find that PTBP1 controls transcript-specific translation. This is consistent with its previously described function in IRES-mediated translation^46^. Whether its differential mode of action on target mRNA is defined by a specific class of motifs that PTBP1 binds to, its subcellular location, or its interaction partners, remains to be resolved. Of note, PTBP1 KO T cells also displayed reduced puromycin incorporation, indicating that PTBP1 regulates global translation as well, possibly by promoting ribosome biogenesis, as recently reported for hematopoietic cells^47^. Combined, these observations indicate a multifactorial activity of PTBP1 on its target mRNA and on translation regulation, with its dominant mode of action possibly determined by the cellular context. It is noteworthy that several predicted PTBP1 binding motifs by SONAR are also binding hubs for other RBPs, including PTBP2, SRSF3, or ELAVL2^36^. However, none of these RBPs were detected in polysome-dense fractions (**Suppl Table8**), suggesting limited effects by these proteins in promoting active translation. Only SRSF3 was detected in the monosome fraction, which could hint to its activity in translation initiation, supported by the previously reported interaction of SRSF3 with the translation initiation factor eIF4A1^48^.

We also followed up on two other RBPs, i.e. HNRNPC and SRSF1, which increased their polysome association in activated T cells. HNRNPC interacts with U-rich elements in the CDS and the 3’UTR in an m6A-dependent manner^49^, and was reported to regulate splicing of translation factors and thus affecting global translation rates in Jurkat cells^50^. Intriguingly, HNRNPC regulates translation of ribosomal proteins and translation factors independently of its splicing activity^51^, indicating activity of this RBP beyond mRNA processing. As HNRNPC alike, also SRSF1 was found in polysomes fractions in mitotic cells^51,52^. SRSF1 regulates the translation of >500 transcripts in mitotic cells, including cell cycle-, transcription- and RNA metabolism-related genes^52^. SRSF1 interacts with purine-rich elements in the CDS and 5’UTR and regulates both alternative splicing and translation of its target mRNAs^52^. It is therefore conceivable that increased polysome association of SRSF1 in activated T cells couples transcript isoform choice with protein output, and that HNRNPC association rather points to a switch from its function as splicing factor towards a translation regulator. Interestingly, even though HNRNPC and SRSF1 depletion led to poor T cell viability and total translation rates, viable KO T cells expressed more effector molecules. This finding could point to a dual function of HNRNPC and SRSF1, by constraining translation of effector molecules as reported for SRSF1-deficient mouse CD8^+^ T cells^53^, yet supporting translation of transcripts required for cellular fitness and homeostasis.

SONAR predicted other interesting putative RBP binding sites regulating polysome occupancy. This included the m6A reader YTHDC1, which was also found with high feature importance in the 3’UTR. YTHDC1 is primarily located within the nucleus where it regulates alternative splicing^54^, and it can act as nuclear export factor^55^. Here, we report that YTHDC1 is enriched in polysome-light fractions of activated T cells, pointing to its putative regulatory activity beyond its currently known activities. SONAR also identified binding hubs for TIA1/TIAL1. TIA1/TIAL1 can interact with U-rich elements (TTTTTC) within 3′UTRs and regulate translation of transcription factors to maintain quiescence in murine naive T cells^27,56^. Interestingly, SONAR and the GFP reporter assay both identified several previously uncharacterized repressive elements, including the GTTTTTC motif. Remarkably, this motif was even more repressive than the well-studied, highly suppressive AU-rich elements. Whether TIA1/TIAL or other U-rich binding RBPs exert the observed strong repressive activity through the GTTTTTC element remains, however, to be determined.

Together, our data support a model in which RBPs act as modulators of translational selectivity and proteome remodeling during T-cell activation. Future work defining their direct mRNA targets, and dissecting their dynamic regulation in activated T cells, will be instrumental for clarifying how RBPs coordinate survival, proliferation and effector function in T cells.

## Materials and Methods

### T cell activation and culture

Human T cells from anonymized healthy donors were used in accordance with the Declaration of Helsinki (Seventh Revision, 2013) after written informed consent (Sanquin). Cryopreserved lymphocytes from buffy coats or elutriation from peripheral blood mononuclear cells (PBMCs) were defrosted, and CD8^+^ T cells were purified with CD8 Microbeads (Miltenyi Biotec) for RNAseq/MS analysis, and through negative selection with CD4 microbeads for follow up experiments according to the manufacturer’s protocol. T cells from pooled PBMCs from multiple donors were used for RNAseq/MS analysis (n=3 pools of 40 donors) and MPRA screen (n=2 pools of 5 donors). T cells were activated with α-CD3/α-CD28 as previously described^2^. Briefly, 24-well plates precoated overnight at 4°C with 4 μg/mL rat α-mouse IgG2a (MW1483, Sanquin) in phosphate-buffered saline (PBS) were washed and coated for >3 h at 37°C with 1 μg/mL α-CD3 (HIT3a, Biolegend). 1×10^6^ CD8^+^ T cells/well were seeded with soluble 1 μg/mL α-CD28 (CD28.2, Biolegend) in 1 mL culture medium (CM: Iscove’s Modified Dulbecco’s Media (IMDM) (LONZA), 100 U/mL penicillin, 100 μg/mL streptomycin, and 2 mM L-glutamine, supplemented with 10% fetal bovine serum (FBS). After 48 h activation at 37°C, 5% CO_2_, T cells were harvested and cultured at a density of 0.8 x 10^6^/mL in standing T25/75 tissue culture flasks (Thermo Scientific) in CM supplemented with 5% heat-inactivated human serum (Sanquin), 5% FBS and 100 IU/mL recombinant human (rh) IL-2 (Proleukin, Clinigen). Medium was refreshed after 2 days. At day 5 of rest, T cells were collected, spun for 8 min at 255 x g (1,200 rpm), and 100 x 10^6^ CD8^+^ T cells per condition were activated for 2 h with 10 ng/ml PMA and 1 μM Ionomycin (Sigma-Aldrich), or T cells were left nonactivated.

### Polysome fractionation for RNA sequencing and MS analysis

Activated and nonactivated T cells were washed twice with ice-cold PBS containing 100 μg/mL cycloheximide (Sigma-Aldrich) and lysed in lysis buffer (110 mM potassium acetate, 20 mM magnesium acetate, 10 mM HEPES pH 7.6, 100 mM potassium chloride, 10 mM magnesium chloride, 0.1% NP-40 and freshly added 2 mM dithiothreitol (DTT), 40 U/mL RNase OUT (both Invitrogen), 1% EDTA-free protease/phosphatase inhibitor cocktail (Thermo Scientific) and 100 μg/ml cycloheximide). For transcriptome and proteome analysis of the full T cell lysate (TCL), 1 x 10^6^ and 2 x 10^6^ T cells from the identical samples were collected, respectively, washed with PBS, and pellets were stored at –80°C. For polysome fractionation, T cell lysates were layered on a 17-50% sucrose gradient (sucrose dilutions in 110 mM potassium acetate, 20 mM magnesium acetate, 10 mM HEPES pH 7.6 and 100 mg/ml cycloheximide) and centrifuged for 140 min at 178,305 x g (38,000 rpm) at 4°C in a SW41Ti rotor. 15 fractions of 750 µl each were harvested, which were divided into equal parts for individual RNA and protein isolation. To isolate RNA from each fraction, samples were treated with 75 μg molecular grade proteinase K (Life Technologies), 1% SDS and 10mM EDTA for 30min at 42°C prior the Phenol/Chloroform extraction. RNA content and RNA integrity of each fraction was determined with the RNA 6000 Nano assay on the Bioanalyser 2100 (Agilent). Fractions were combined into 4 pools as indicated in Fig 2B: free RNA (fractions #1-3), monosome-bound RNA (#4-5), polysome light (#6-10) and polysome heavy-bound RNA (#11-15). Fractions were then purified and concentrated with RNA Clean & Concentrator-5 kit (Zymo Research). RNA for the whole cell transcriptome analysis was isolated with Quick-RNA Microprep kit (Zymo Research).

Proteins were precipitated from each fraction pool with ethanol. SDS was first added to samples to a final concentration of 1%. One volume of pooled fractions was mixed with 9 volumes of 100% ice-cold ethanol and kept at –20°C overnight. Samples were spun at 16,100 x g (13,200 rpm) for 10 min at 4°C and protein pellets were further washed with 80% ice-cold ethanol, spun and air dried to eliminate any ethanol residue. Protein pellets were kept at –80°C until further processing for MS.

### Sample preparation for RNA-sequencing

Libraries were prepared with the NEBNext Ultra II Directional RNA Library Prep Kit for Illumina including rRNA depletion with NEBNext rRNA depletion kit (according to manufacturer’s protocol). Libraries were not size selected. Sequencing was performed using the NovaSeq6000 (Illumina), at a target depth of 30 million reads for polysome fractions, and 60 million reads for full transcriptome, all paired-end 150 nt reads (GenomeScan, Leiden).

### RNA-sequencing analysis

RNA-sequencing reads were quasi-mapped on the human transcriptome GRCh38 from Gencode (version 38, October 2021)^57^ using Salmon tool version 1.5.2^58^. An average of 33×10^6^ reads for each sample was mapped to the transcriptome. Due to the low number of mapped reads (8.2×10^6^; 16.6% of total reads, caused by a few overrepresented sequences), the sample of nonactivated, monosome fraction (pool 2) was excluded from further analysis. Mapping results were summarized on a gene-level using tximport in R^59^. DESeq2 version 1.34.0 was used for differential expression analysis^60^. A gene was considered differentially expressed if Benjamini-Hochberg adjusted *p*-value was >0.001, absolute value of log2 fold change >1, and the mean of normalized counts across samples at least 10.

### RNA level estimation

RNA-sequencing reads were mapped to human genome GRCh38 primary assembly from Gencode (version 38, December 2021)^57^ using STAR version 2.7.9a^61^ to count all reads that originated from a given gene, including the reads originating from intronic regions. On average, 67% of reads were uniquely mapped to the genome. For each gene, reads per kilobase (RPK) were calculated by dividing read count by gene length. Per million scaling factor was calculated as the sum of all RPKs divided by 10^6^. TPMs were then calculated as corresponding RPKs divided by the scaling factor. Mass of RNA was calculated as *m* = TPM × *M* / *N*_A_, where molar mass was estimated based on RNA length as *N*(nucleotides) × 320.5 + 159 (approximate molecular weight of a nucleotide and addition of 159 to take into account the molecular weight of a 5’triphosphate). For RNAs that contain multiple isoforms, an average length was calculated either by assuming equal amount of each isoform, or by taking the proportion of each isoform into account (when they were present in the output of Salmon quasi-mapping). Masses were then scaled to the total expected mass, which was calculated as RNA mass measured after PCR amplification, multiplied by the percentage of mapped reads. The mass of each RNA species in the initial sample was calculated based on known sample dilutions and the number of PCR cycles (Supplemental Figure 2A). The number of RNA molecules was then calculated as *N* = *m* × *N*_A_ / *M*.

To describe the RNA distribution between fractions, a ratio of ribosome-bound:free RNA (RF) was calculated for each gene separately for non-activated and activated state. To capture the difference in RNA distribution, the two RF ratios for each RNA were divided (RF_PMA_/RF_rest_); a value above 1 indicates a shift from free to ribosome-bound RNA upon activation.

### Clustering RNA distribution profiles

To obtain the distribution profile of an RNA across the four fraction pools (free RNA, monosomes, polysomes light, and polysomes heavy), the estimated RNA amount in each fraction was divided by the total amount of the same RNA across all fractions, thereby obtaining RNA percentage in each fraction. For clustering, we took 1) each RNA that is differentially expressed in at least one of the polysome fractions and 2) all non-differentially expressed RNAs from all fractions as long as their mean TPM across replicates was >0.1 in both resting and activated state. Distribution profiles of these RNAs were clustered by hierarchical clustering with Ward’s linkage method (“ward.D” in R”), using *stats* package version 4.1.2 in R. Pearson’s correlation subtracted from 1 was used as a distance measure. R implementation of *G:profiler*^62^ was applied to determine significantly enriched (*p*-value < 0.05) gene ontology (GO) biological process terms within each of the 5 obtained clusters. Because clustering was performed in a non-biased manner, i.e., did not take into account whether RNA distribution profiles stemmed from nonactivated or activated RNA distribution profiles, GO enrichment analysis was also performed for genes displaying resting-to-activated shifts between clusters.

### Machine learning classification

#### Sequence feature generation

The features library for XGBoost was built as described previously^13^. Briefly, the human transcriptome was searched for RBP binding motifs extracted from ATtRACT database (*Homo sapiens)*^36^, found within the coding sequence (CDS) and untranslated regions (5’-and 3’-UTR) as separate entities. For RF models, additional 552 5-mer RBP-interacting motifs identified by Dominguez *et al.*^63^ were included, resulting in a total of 7,488 features. Furthermore, it included GC content, sequence length, codon usage (normalized for the length of CDS), and selected mRNA modifications (m6A, m6C, A-to-I editing, m1A, m7G) that were obtained from RMVar database^64^. When multiple annotated isoforms were present, each feature was averaged across isoforms to obtain a gene-based feature library.

### Modelling

XGBoost (v. 1.4.1.1)^65^ in *Caret* (v. 6.0-90) was used to identify sequence determinants that explain changes in RNA distribution profiles over polysome fractions. The data were split prior to training into 80 or 70% for training, and 20 or 30% held-out test set for performance testing. Due to class imbalances, down sampling was used. Non-informative features were excluded from model training using *nearZeroVar* function from *Caret*. Each model was first trained using a coarse grid of parameters (nrounds, max_depth, colsample_bytree, eta, gamma, min_child_weight, subsample) and a single repeat of a 3-fold cross-validation, followed by a finer grid and three repeats of a 10-fold cross-validation. We trained two prediction models, each performing a binary classification: 1) DEG with a RF_PMA_/RF_rest_ ratio of >1 vs. <1, 2) nonDEG with RF_PMA_/RF_rest_ ratio of >1 vs. <1. DEG and nonDEG were determined based on DE analysis (DESeq2) of TCL samples, where nonDEG were filtered by mean TPM value across replicates. Cut-off for analysis inclusion was mean value > 0.1 in both nonactivated and activated state.

Finally, the importance of RNA features was assessed. Features were ranked and compared to each other (scale 0-100, where 0 is assigned to a feature that was not useful in decision trees, and 100 to the most useful feature), to uncover the most useful RNA sequence features for classification with XGBoost.

### Sample preparation for MS

Reduction of disulphide bridges in cysteine containing proteins was performed with DTT (56°C, 30 min, 10 mM in 50 mM HEPES, pH 8.5). Reduced cysteines were alkylated with 20 mM 2-chloroacetamide (30 min at room temperature in 50 mM HEPES, pH 8.5). Samples were prepared using the SP3 protocol^66,67^. Trypsin (sequencing grade, Promega) was added in an enzyme to protein ratio 1:50 for overnight digestion at 37°C. Peptides were recovered in HEPES buffer by collecting supernatant on magnet and combining with a second elution wash of beads with HEPES buffer. Peptides were labelled with TMT6plex Isobaric Label Reagent (ThermoFisher) according to the manufacturer’s instructions. Samples were combined for the TMT6plex and for further sample clean up with an OASIS® HLB mElution Plate (Waters). Offline high pH reverse phase fractionation was carried out on an Agilent 1200 Infinity high-performance liquid chromatography system, equipped with a Gemini C18 column (3 mm, 110 Å, 100 x 1.0 mm, Phenomenex).

An UltiMate 3000 RSLC nano LC system (Dionex) was fitted with a trapping cartridge (µ-Precolumn C18 PepMap 100, 5µm, 300 µm i.d. x 5 mm, 100 Å) and an analytical column (nanoEase M/Z HSS T3 column 75 µm x 250 mm C18, 1.8 µm, 100 Å, Waters). Trapping was carried out with a constant flow of trapping solution (0.05% trifluoroacetic acid in water) at 30 µL/min onto the trapping column for 6 min. Peptides were eluted via the analytical column running solvent A (0.1% formic acid in water) with a constant flow of 0.3 µL/min, with increasing percentage of solvent B (0.1% formic acid in acetonitrile). The outlet of the analytical column was coupled directly to an Orbitrap QExactive plus Mass Spectrometer (Thermo) using the Nanospray Flex ion source in positive ion mode.

The peptides were introduced into the QExactive plus via a Pico-Tip Emitter 360 µm OD x 20 µm ID; 10 µm tip (New Objective) and an applied spray voltage of 2.2 kV. The capillary temperature was set at 275°C. Full mass scan was acquired with mass range 375-1200 m/z in profile mode with resolution of 70000. The filling time was set at maximum of 100 ms with a limitation of 3×10^6^ ions. Data dependent acquisition (DDA) was performed with the resolution of the Orbitrap set to 17500, with a fill time of 50 ms and a limitation of 2×10^5^ ions.

A normalized collision energy of 32 was applied. Dynamic exclusion time of 20 s was used. The peptide match algorithm was set to ‘preferred’ and charge exclusion ‘unassigned’, charge states 1,5 - 8 were excluded. MS2 data was acquired in profile mode.

### MS proteomics data processing

IsobarQuant^68^ and Mascot (v2.2.07) were used to process the acquired data, which was searched against the Uniprot Homo sapiens proteome database (UP000005640) containing common contaminants and reversed sequences. The following modifications were included into the search parameters: Carbamidomethyl (C) and TMT6 (K) (fixed modification), Acetyl (Protein N-term), Oxidation (M) and TMT6 (N-term) (variable modifications). For the full scan (MS1) a mass error tolerance of 10 ppm and for MS/MS (MS2) spectra of 0.02 Da was set. Further parameters were set: trypsin as protease with an allowance of maximum two missed cleavages; a minimum peptide length of seven amino acids; at least two unique peptides were required for protein identification. The false discovery rate on peptide and protein level was set to 0.01.

### MS proteomics analysis

DEP package version 1.14.0 was applied for differential protein abundance analysis^69^ using nonactivated-activated sample pairs. Proteins with any missing values across the triplicate samples were filtered out to avoid imputation of missing values. A protein was considered differentially expressed (DE) if adjusted *p*-value was below 0.05 and the log2 fold change >1.5 (this threshold was used due to ratio compression with TMT-labelling). RNA-binding proteins were annotated using a previously published RBP list^29^.

### GFP reporters

As a backbone, a control sequence of 166 nt was designed lacking all 6-12 mer motifs that displayed any importance >0 in any of the ML models, as well as motifs of length 4-5 nt with feature importance above 4. Furthermore, the RNAfold webserver^70^ was used to exclude putative formation of strong secondary structure elements in the scrambled sequence. Each motif was introduced 6 times into 3’UTRs of a GFP reporter, with distances between the motif repeats of 12, 12, 25, 26 and 33 nucleotides, respectively, and cloned into BamHI and NotI sites of pRETRO-SUPER GFP downstream of GFP coding sequence.

### Virus production

FLYRD18 retroviral packaging cells [European Collection of Authenticated Cell Cultures (ECACC) 95091902] were cultured in Iscove’s Modified Dulbecco’s Media (IMDM) (LONZA) supplemented with 10% FBS, 100 U/mL penicillin, 100 μg/mL streptomycin, and 2 mM L-glutamine at 37°C, 5% CO_2_. Cells were plated at a density of 1,2 × 10^5^ per well in a 6-well plate for 16 hours prior to transfection with GeneJammer (Agilent) and kept at 32°C, 5% CO_2_. The retroviral supernatant was harvested after 48 hours and was either used freshly or was snap-frozen in liquid nitrogen and stored at −80°C until further use.

### Retroviral transduction

Transduction of T cells was performed with Retronectin (Takara) as previously described^2^. Briefly, non-tissue cultured treated 24-well plates were coated overnight with 10 μg/mL Retronectin (Takara), washed once with PBS prior to adding viral supernatant. Plates were centrifuged for 30 min at 4**°**C at 4500 rpm (2820g). 0,7×10^6^ T cells were added/well, spun for 5 min at 1000 rpm (180 g), and incubated overnight at 37**°**C. The following day, cells were harvested and cultured in T25/75 flasks at a concentration of 0.8×10^6^ cells/mL for 6-8 days in presence of rhIL-2 and rhIL-15.

### Genetic modification of T cells with Cas9 RNPs

gRNAs were designed in Benchling (https://benchling.com). Cas9 RNP production and T cell nucleofection was performed as previously described^2^. Briefly, Alt-R crRNA were reconstituted to 100 μM in Nuclease Free Duplex buffer (all Integrated DNA Technologies) and mixed at equimolar ratio with tracrRNA (i.e. 4.5 μL total crRNA + 4.5 μL tracrRNA) in nuclease-free PCR tubes and denatured by heating at 95°C for 5 min in a thermocycler. Alternatively, 4.5 μL of Alt-R sgRNA were used. As a negative control, nontargeting negative control crRNA #1 was used (Integrated DNA Technologies). Nucleic acids were cooled down to room temperature prior to mixing them with 30 μg Alt-R™ S.p. Cas9 Nuclease V3 (IDT) to produce Cas9 ribonuclear proteins (RNPs). Mixture was incubated at room temperature for at least 10 min prior to nucleofection. For nucleofection, human CD3^+^ T cells were activated for 72 h with α-CD3/α-CD28. Cells were electroporated in 16-well strips in a 4D Nucleofector X unit with program EH100 and P2 buffer (Lonza). Right after nucleofection cells were refreshed and kept in IMDM supplemented with 5% heat-inactivated human serum, 5% FBS and 100 IU/mL recombinant human (rh) IL-2 and 10 ng/mL rhIL-15 (Peprotech). Knockout efficiency was determined on day 7-10 after electroporation by Western blot or flow cytometry.

### Flow cytometry and intracellular cytokine staining

T cells were washed with FACS buffer (PBS, containing 1% FBS and 2 mM EDTA) and labeled for 20 min at 4 °C with α-CD4 (SK3 and RPA-T4), α-CD8 (SK1, both Biolegend), α-IFN-g (4S.B3, eBioscience), α-Puromycin (12D10, Merck), α-Granzyme B (GB11), α-CD107a (H4A3, both BD Biosciences), α-IL-2 (MQ1-17H12), and α-TNF (MAb11), α-Perforin (BD-48), α-CD25 (BC96), α-CD69 (FN50), α-CD137 (4B4-1, all Biolegend). Dead cells were excluded with Near-IR (Life Technologies). For Puromycin incorporation assay, cells were treated with 5 μg/mL Puromycin dihydrochloride (Merck) for the last 10min of culture, washed twice with PBS, stained intracellularly and washed twice prior to acquisition. For intracellular cytokine staining, cells were cultured with 1 μg/mL brefeldin A and 2 μM Monensin from the start of assay, fixed and permeabilized with Cytofix/Cytoperm kit (BD Biosciences) prior to acquisition using FACSymphony or LSRFortessa. Data were analyzed with FlowJo (BD Biosciences, version 10).

### MPRA screen

The retroviral library was produced by transfecting 4.5 μg of the library plasmid DNA into FLYRD18 retroviral packaging cells, as described above. CD8^+^ T cells were isolated from PBMCs from 2 pools of 5 donors, activated with α-CD3/α-CD28 for 48h and transduced as described above. 24h after transduction, cells were washed and nucleofected with Cas9 RNPs, as described above. Transduction efficiencies were kept between 8-12% to minimize multiple transgene integrations. After expansion for 7 days, T cells were harvested and stained with Near-IR live/dead marker (Life Technologies) for dead cell exclusion, in addition to α-CD8 and α-CD4. The top and bottom 15% GFP-expressing CD8^+^ T cells, in addition to the full GFP-positive fraction, were sorted on an Aria II (BD Biosciences). Fluorescence-activated cell sorted cells were subsequently pelleted and resuspended in 1 μl of DirectPCR Lysis Reagent (Viagen) per 5,000 cells, supplemented with proteinase K (0.5 mg/ml; Viagen). The obtained lysate was incubated at 55°C for 3 hours, followed by 85°C for 45 min and 95°C for 5 min.

### 3′UTR reporter assay library amplification and sequencing

An initial preamplification of the synthetic 3′UTRs was performed on 10 μl of lysate (±50,000 cells) in a 50-μl reaction volume using NEBnext master mix (NEB) and 10 μM forward (5′-AAAGACCCCAACGAGAAGC-3′) and reverse (5′-AGTCTATAGCTACTAGGCG-3′) primers. This PCR amplifies three (out of five) of the motif sequences. An initial touchdown stage was performed by reducing the annealing temperature 1°C per cycle from 58° to 50°C (eight cycles), followed by six cycles at 50°C. Extension time was kept at 20 s. PCR products were purified using Cytiva Sera-Mag Select at a 1.5:1 Sera-Mag:PCR-product ratio, following the manufacturer’s protocol. Each of these samples were subsequently used as template for a second PCR, with (forward: 5′-ACACTCTTTCCCTACACGACGCTCTTCCGATCTAGACCCCAACGAGAAGCGCGATCAC-3′, reverse: 5′-GACTGGAGTTCAGACGTGTGCTCTTCCGATCTAGTCTATAGCTACTAGGCGATA-3′) primers to amplify the product and attach partial Illumina adapters. This second PCR was performed using NEBnext master mix and consisted of 10-15 cycles with a constant annealing temperature of 65°C and extension time of 30 s. Correct products (±275 base pairs) were purified using Cytiva Sera-Mag Select at a 1.5:1 Sera-Mag:PCR-product ratio, following the manufacturer’s protocol. Purified products were indexed with custom Unique Dual Index, pooled, purified, and sequenced at the genomics core facility of the Netherlands Cancer institute (paired-end sequencing, 300 cycles). Raw data are available at GEO (see below).

### 3′UTR parallel reporter analysis

The obtained read 2 data contained the three motif sequences. First, the sequencing reads were trimmed using cutadapt to remove constant sequences and PCR handles. Second, trimmed reads were aligned to the synthetic UTR library using bowtie2, which was performed using the ‘local’ alignment and ‘very-sensitive’ options. Third, aligned reads were quantified using htseq-count, using a custom gtf file constructed from the synthetic UTR library. Motifs with less than 20 read counts across all samples were filtered out, and library size of each sample was scaled to 10,000 to allow comparison between samples.

## Data and code availability

The MS data of the polysome fractions has been deposited to the ProteomeXchange Consortium via the PRIDE partner repository with the dataset identifier PXD029199. The RNAseq data have been deposited to ENA European Nucleotide Archive with the project accession PRJEB111352. The MPRA data have been deposited to DANS Data Station Life Sciences and are available at https://doi.org/10.17026/LS/MCQNZ2. Codes used in the analysis described in this study are available at https://github.com/nikolinasostaric/CD8_polysome.

## Statistical analysis

Statistical analysis and visualization were performed using R or GraphPad Prism (version 10.1.0). Sample sizes, statistical tests and cut offs for significance are detailed in the respective figure legends.

## Acknowledgments

We thank K. Rooijers, S. Tol and E. Mul for technical help. This research was supported by European Research council (ERC) consolidator award PRINTERS 817533 and Oncode Institute, both to M.C.W., and Landsteiner Foundation for Blood Transfusion (LSBR) Research grant 2103 to M.C.W. and B.P.

## Author contributions

B.P., B.P.N. and M.C.W. conceived and designed experiments. B.P., K.B., A.B., S.E., A.G. and M.R. performed experiments. B.P., B.P.N., N.Š., K.B., N.K. and F.S. analyzed the data. J.I.P-P. provided intellectual input. B.P. and M.C.W. wrote the manuscript. All authors have read and approved the submitted version. M.C.W. provided supervision and funding.

## Competing interests

The authors declare no competing interests.

---

**Supplementary Figure 1.**
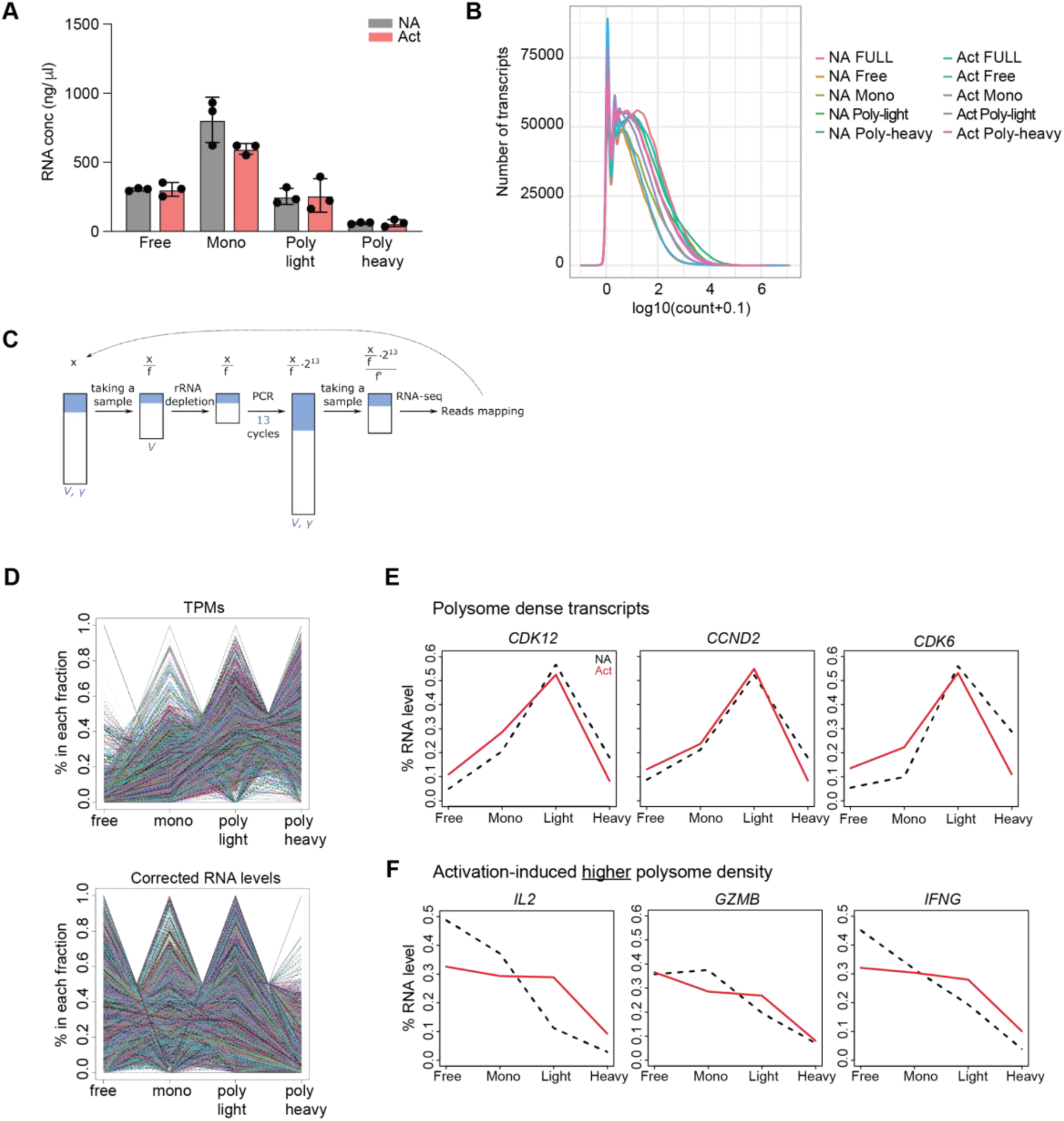
A) Concentrations of RNA isolated from each polysome fraction (see Figure 2B). Data are presented as mean ± SD. B) Transcript count distributions across different conditions (log 10 counts+0.1, weighted by replicates). C) Corrected levels of RNA isolated from fractionated cell lysates was estimated based on RNA-seq data and quantitative values from the sample preparation steps (denoted in blue). In the scheme, an amount of a theoretical mRNA is denoted with x, which is then followed upon different steps. D) RNA distribution profiles across the fractions for all genes, calculated with either TPMs (up) or corrected RNA levels (down). E-F) RNA distribution plots for indicated genes in nonactivated (NA) and activated (Act) CD8^+^ T cells.

**Supplementary Figure 2.**
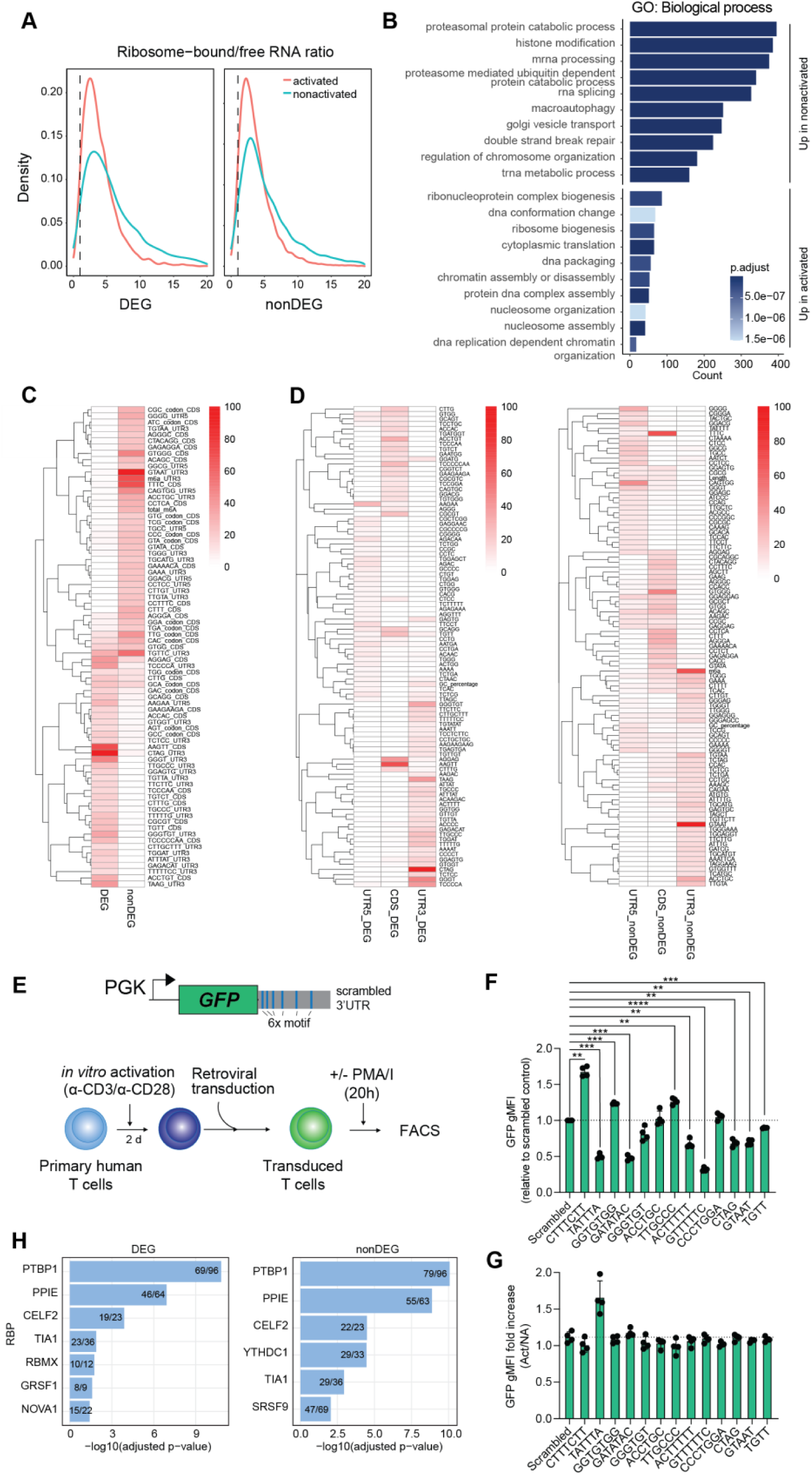
Sequence features predict RNA shifts between fractions. A) Ribosome bound vs. free RNA ratios among DEG and nonDEGs between activated and nonactivated T cells. B) GO enrichment analysis of biological processes within groups of genes that show shift towards higher ribosomal occupancy depending on the T cell activation status. C) Heatmaps showing the importance of the top predictive features from ribosome-bound/free (RF) RNA models between the DEG and nonDEG models and D) between different RNA regions in DEG and nonDEG. E) Experimental setup to test the effect of sequences with high feature importance in the 3’TUR, as defined by SONAR models. Selected motifs were introduced 6 times into the 3’UTR of the GFP reporter constructs. Primary human T cells were retrovirally transduced with GFP constructs and GFP was measured by FACS after resting and reactivation for 20 h with PMA/I. F) GFP gMFI in transduced resting human CD8^+^ T cells. Data depict mean ± SD of 4 donors, representative of at least 2 independently performed experiments (Repeated measures one-way ANOVA with Dunnett’s multiple comparison to the control; **P < 0.01, ***P < 0.001; ****P < 0.0001.). G) Fold increase in GFP gMFI levels upon activation with PMA/I (Act), compared to nonactivated cells (NA). H) Enrichment of RBP binding sites with importance >0 in 3’UTR in RF model. Fischer’s exact test. Graphs show RBPs with adjusted p-value < 0.05.

**Supplementary Figure 3.**
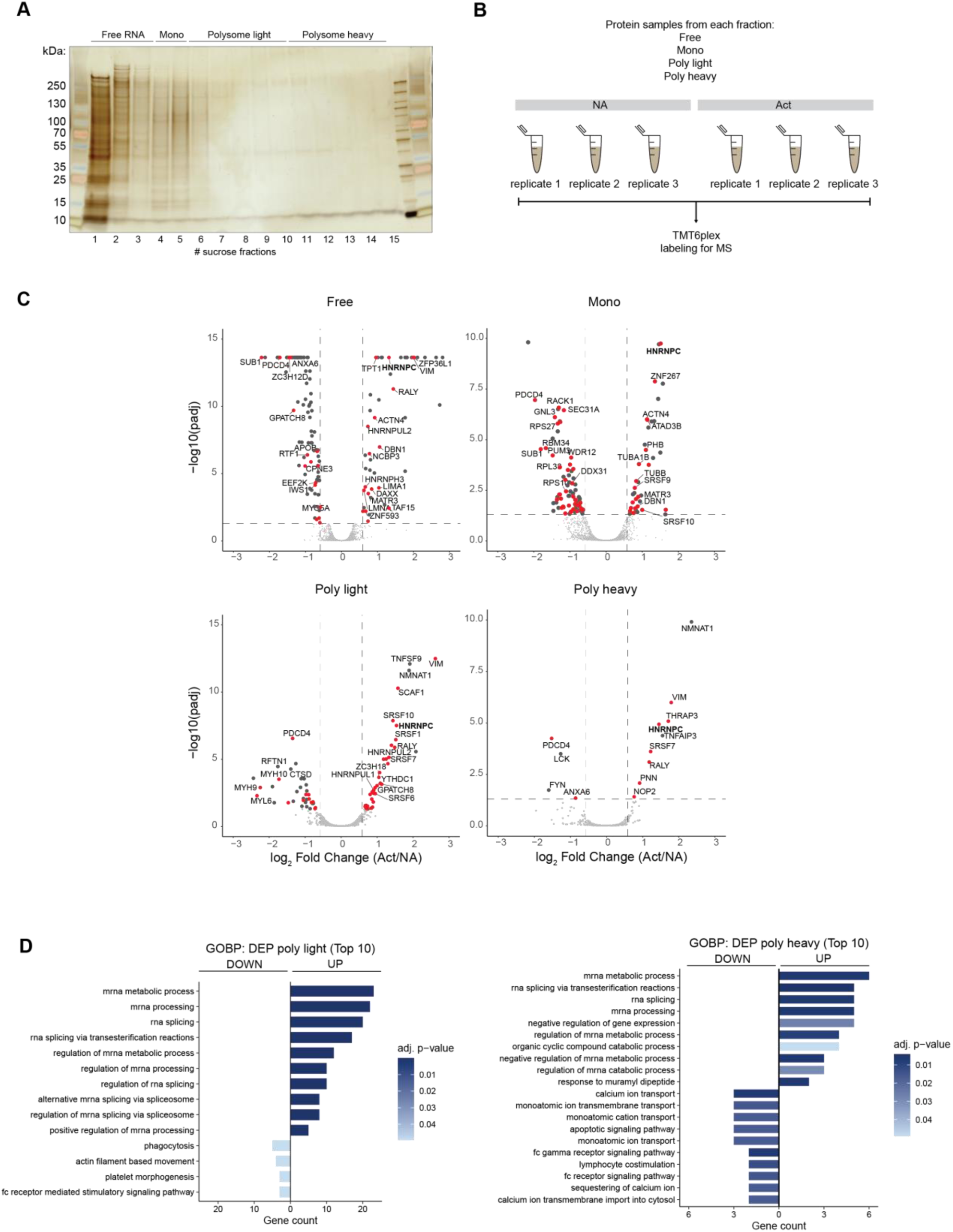
A) Silver staining to measure proteins isolated from polysome fractions. Proteins from 15 fractions were pooled as indicated in the figure and subjected to MS analysis. B) Schematic overview of protein samples pooling for TMT6 labeling and MS analysis. C) Volcano plots showing differentially expressed proteins in polysome fractions from activated (Act) vs. nonactivated (NA) T cells. RBPs are indicated as red dots. D) GO enrichment analysis of biological processes within DEP in polysome light and heavy fractions upon T cell activation.

**Supplementary Figure 4.**
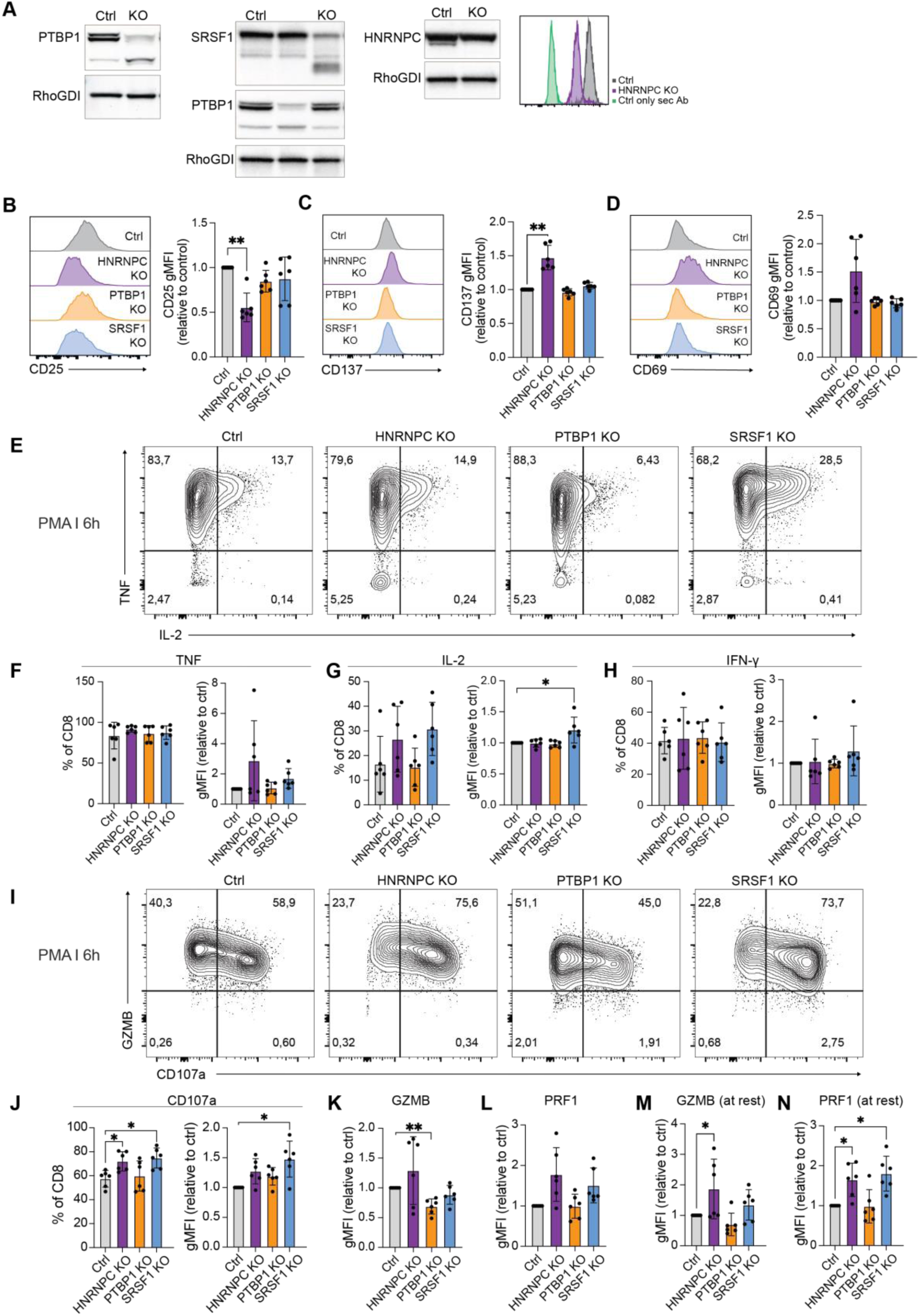
A) Representative immunoblots and histogram of human CD8^+^ T cells 7-9 days after CRISPR/Cas9 gene-editing for indicated RBP, or with non-targeting crRNA. B-D) Representative histograms and fold change of CD25 (B), CD137 (C) and CD69 (D) gMFI levels in RBP KO CD8^+^ T cells compared to controls. Compiled data of 6 donors from 2 independently performed experiments. Data are presented as mean ± SD. Repeated measures one-way ANOVA with Dunnett’s multiple comparison to the control and unpaired t test (**P<0.05). E) Representative plots of TNF and IL-2 expression in RBP KO CD8^+^ T cells and control at 6h after activation with PMA/I. F-H) Percentage and gMFI of TNF (H), IL-2 (I), and IFN-γ (J) producing CD8^+^ T cells at 6h after activation with PMA/I. Data are presented as mean ± SD of 6 donors, compiled data from 2 independently performed experiments. Repeated measures one-way ANOVA with Dunnett’s multiple comparison to the control and unpaired t test (*P<0.05). I) Representative plots of Granzyme B and CD107a expression in RBP KO CD8^+^ T cells and control at 6h after activation with PMA/I. J-L)Percentage and gMFI of CD107a (J), granzyme B (K) and perforin (L) in CD8^+^ T cells at 6h after activation with PMA/I. M-N) gMFI of granzyme B (M) and perforin (N) in CD8^+^ T cells at rest. Data are presented as mean ± SD of 6 donors, compiled from 2 independently performed experiments. Repeated measures one-way ANOVA with Dunnett’s multiple comparison to the control and unpaired t test (*P<0.05, **P<0.05).

**Supplementary Figure 5.**
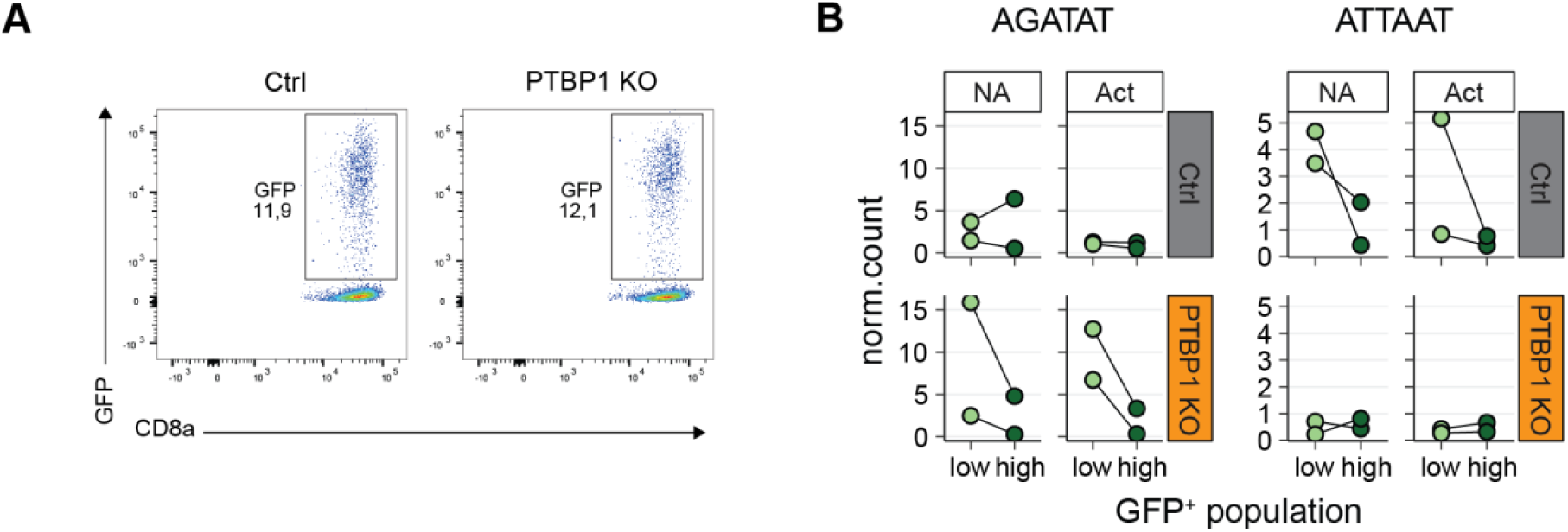
A) Representative GFP expression in PTBP1 KO and control CD8^+^ T cells that were transduced with the MPRA library of synthetic 3’UTRs combined with GFP reporter gene (Fig 6A). B) Graphs depict normalized counts of indicated motifs in GFP^hi^ and GFP^low^ sorted CD8^+^ T cells.

## Notes

### Competing Interest Statement

The authors have declared no competing interest.

## References

1. Han, Q. et al. Polyfunctional responses by human T cells result from sequential release of cytokines. Proceedings of the National Academy of Sciences 109, 1607–1612 (2012).

2. Popović, B. et al. Time-dependent regulation of cytokine production by RNA binding proteins defines T cell effector function. Cell Rep. 42, 112419 (2023).

3. Nicolet, B. P., Guislain, A. & Wolkers, M. C. Combined Single-Cell Measurement of Cytokine mRNA and Protein Identifies T Cells with Persistent Effector Function. J Immunol 198, 962–970 (2017).

4. Jurgens, A. P., Popović, B. & Wolkers, M. C. T cells at work: How post-transcriptional mechanisms control T cell homeostasis and activation. Eur. J. Immunol. 51, 2178–2187 (2021).

5. Salerno, F. et al. Translational repression of pre-formed cytokine-encoding mRNA prevents chronic activation of memory T cells. Nat. Immunol. 19, 828–837 (2018).

6. Wolf, T. et al. Dynamics in protein translation sustaining T cell preparedness. Nat Immunol 21, 927–937 (2020).

7. Tavernier, S. J. et al. A human immune dysregulation syndrome characterized by severe hyperinflammation with a homozygous nonsense Roquin-1 mutation. Nat. Commun. 10, 4779 (2019).

8. Aichele, P. et al. Immunopathology caused by impaired CD8 ^+^ T-cell responses. Eur. J. Immunol. 52, 1390–1395 (2022).

9. Dai, Y.-T. et al. Transcriptome-wide subtyping of pediatric and adult T cell acute lymphoblastic leukemia in an international study of 707 cases. Proceedings of the National Academy of Sciences 119, (2022).

10. Salerno, F., Turner, M. & Wolkers, M. C. Dynamic Post-Transcriptional Events Governing CD8+ T Cell Homeostasis and Effector Function. Trends Immunol. 41, 240–254 (2020).

11. Turner, M. & Petkau, G. RNA-binding proteins and ribonucleoproteins as determinants of immunity. Nat. Rev. Immunol. https://doi.org/10.1038/s41577-025-01254-2 (2026) doi:10.1038/s41577-025-01254-2.

12. de Sousa Abreu, R., Penalva, L. O., Marcotte, E. M. & Vogel, C. Global signatures of protein and mRNA expression levels. Mol. Biosyst. https://doi.org/10.1039/b908315d (2009) doi:10.1039/b908315d.

13. Nicolet, B. P. et al. Learning the sequence code of protein expression in human immune cells. Sci. Adv. 11, (2025).

14. Nicolet, B. P. & Wolkers, M. C. The relationship of mRNA with protein expression in CD8+ T cells associates with gene class and gene characteristics. PLoS One 17, e0276294 (2022).

15. Howden, A. J. M. et al. Quantitative analysis of T cell proteomes and environmental sensors during T cell differentiation. Nat. Immunol. 20, 1542–1554 (2019).

16. Lattanzio, M. V. et al. An early mTOR-dependent window during human T cell activation programs T cell state. Preprint at 10.64898/2026.01.29.702520 (2026).

17. Weerakoon, H. et al. Integrative temporal multi-omics reveals uncoupling of transcriptome and proteome during human T cell activation. NPJ Syst. Biol. Appl. 10, 21 (2024).

18. Nicolet, B. P., Zandhuis, N. D., Lattanzio, V. M. & Wolkers, M. C. Sequence determinants as key regulators in gene expression of T cells. Immunol. Rev. imr.13021 (2021) doi:10.1111/imr.13021.

19. Cho, S., Dong, J. & Lu, L. Cell-intrinsic and -extrinsic roles of miRNAs in regulating T cell immunity. Immunol. Rev. 304, 126–140 (2021).

20. Gebauer, F., Schwarzl, T., Valcárcel, J. & Hentze, M. W. RNA-binding proteins in human genetic disease. Nat. Rev. Genet. 22, 185–198 (2021).

21. Schuschel, K. et al. RNA-Binding Proteins in Acute Leukemias. Int. J. Mol. Sci. 21, 3409 (2020).

22. Moore, M. J. et al. ZFP36 RNA-binding proteins restrain T cell activation and anti-viral immunity. Elife 7, e33057 (2018).

23. Matheson, L. S. et al. Multiomics analysis couples mRNA turnover and translational control of glutamine metabolism to the differentiation of the activated CD4+ T cell. Sci. Rep. 12, 19657 (2022).

24. Essig, K. et al. Roquin targets mRNAs in a 3′-UTR-specific manner by different modes of regulation. Nat. Commun. 9, 3810 (2018).

25. Lingel, H. et al. CTLA-4-mediated posttranslational modifications direct cytotoxic T-lymphocyte differentiation. Cell Death Differ. 24, 1739–1749 (2017).

26. Lattanzio, M. V. et al. Single-molecule imaging of transcription dynamics, RNA localization and fate in human T cells. EMBO J. 44, 6732–6749 (2025).

27. Osma-Garcia, I. C. et al. Post-transcriptional regulation by TIA1 and TIAL1 controls the transcriptional program enforcing T cell quiescence. Preprint at 10.1101/2024.09.03.608755 (2024).

28. Hoefig, K. P. et al. Defining the RBPome of primary T helper cells to elucidate higher-order Roquin-mediated mRNA regulation. Nat. Commun. 12, 5208 (2021).

29. Zandhuis, N. D., Nicolet, B. P. & Wolkers, M. C. RNA-Binding Protein Expression Alters Upon Differentiation of Human B Cells and T Cells. Front. Immunol. 12, 717324 (2021).

30. Lattanzio, M. V. et al. Mapping the dynamic RNA binding proteome in human effector T cells identifies differentiation and cytotoxicity regulators. Preprint at 10.64898/2025.12.05.692493 (2025).

31. Perez-Perri, J. I. et al. Discovery of RNA-binding proteins and characterization of their dynamic responses by enhanced RNA interactome capture. Nat. Commun. 9, 4408 (2018).

32. Tan, H. et al. Integrative Proteomics and Phosphoproteomics Profiling Reveals Dynamic Signaling Networks and Bioenergetics Pathways Underlying T Cell Activation. Immunity 46, 488–503 (2017).

33. Asano, Y. et al. Nuclear polarization to the immune synapse facilitates an early transcriptional burst. Sci. Immunol. 10, (2025).

34. Sinclair, L. V. & Cantrell, D. A. Protein Synthesis and Metabolism in T Cells. Annu. Rev. Immunol. 43, 343–366 (2025).

35. Schwanhäusser, B. et al. Global quantification of mammalian gene expression control. Nature 473, 337–342 (2011).

36. Giudice, G., Sánchez-Cabo, F., Torroja, C. & Lara-Pezzi, E. ATtRACT—a database of RNA-binding proteins and associated motifs. Database 2016, baw035 (2016).

37. Dorrello, N. V. et al. S6K1- and ßTRCP-Mediated Degradation of PDCD4 Promotes Protein Translation and Cell Growth. Science (1979). 314, 467–471 (2006).

38. Benbarche, S. et al. GPATCH8 modulates mutant SF3B1 mis-splicing and pathogenicity in hematologic malignancies. Mol. Cell 84, 1886–1903.e10 (2024).

39. Schmidt, E. K., Clavarino, G., Ceppi, M. & Pierre, P. SUnSET, a nonradioactive method to monitor protein synthesis. Nat. Methods 6, 275–277 (2009).

40. D’Angeli, V. et al. Polypyrimidine tract binding protein 1 regulates the activation of mouse CD8 T cells. Eur. J. Immunol. 52, 1058–1068 (2022).

41. Klein, K. et al. T Cell-Intrinsic CDK6 Is Dispensable for Anti-Viral and Anti-Tumor Responses In Vivo. Front. Immunol. 12, (2021).

42. Lea, N. C. et al. Commitment point during G0-->G1 that controls entry into the cell cycle. Mol. Cell. Biol. 23, 2351–61 (2003).

43. Monzón-Casanova, E. et al. Polypyrimidine tract-binding proteins are essential for B cell development. Elife 9, (2020).

44. Matus-Nicodemos, R. et al. Polypyrimidine Tract-Binding Protein Is Critical for the Turnover and Subcellular Distribution of CD40 Ligand mRNA in CD4+ T Cells. The Journal of Immunology 186, 2164–2171 (2011).

45. La Porta, J., Matus-Nicodemos, R., Valentín-Acevedo, A. & Covey, L. R. The RNA-Binding Protein, Polypyrimidine Tract-Binding Protein 1 (PTBP1) Is a Key Regulator of CD4 T Cell Activation. PLoS One 11, e0158708 (2016).

46. Mitchell, S. A. et al. Identification of a motif that mediates polypyrimidine tract-binding protein-dependent internal ribosome entry. Genes Dev. 19, 1556–1571 (2005).

47. Rehn, M. et al. PTBP1 promotes hematopoietic stem cell maintenance and red blood cell development by ensuring sufficient availability of ribosomal constituents. Cell Rep. 39, 110793 (2022).

48. Kim, J., Park, R. Y., Kee, Y., Jeong, S. & Ohn, T. Splicing factor SRSF3 represses translation of p21cip1/waf1 mRNA. Cell Death Dis. 13, 933 (2022).

49. Liu, N. et al. N6-methyladenosine-dependent RNA structural switches regulate RNA–protein interactions. Nature 518, 560–564 (2015).

50. Dróżdż, M. et al. Immediate early splicing controls translation in activated T-cells and is mediated by hnRNPC2 phosphorylation. EMBO J. 44, 1692–1723 (2025).

51. Aviner, R. et al. Proteomic analysis of polyribosomes identifies splicing factors as potential regulators of translation during mitosis. Nucleic Acids Res. 45, 5945–5957 (2017).

52. Maslon, M. M., Heras, S. R., Bellora, N., Eyras, E. & Cáceres, J. F. The translational landscape of the splicing factor SRSF1 and its role in mitosis. Elife 3, (2014).

53. Zhu, G.-Q. et al. Targeting SRSF1 improves cancer immunotherapy by dually acting on CD8+T and tumor cells. Signal Transduct. Target. Ther. 10, 25 (2025).

54. Xiao, W. et al. Nuclear m6A Reader YTHDC1 Regulates mRNA Splicing. Mol. Cell 61, 507–519 (2016).

55. Roundtree, I. A. et al. YTHDC1 mediates nuclear export of N6-methyladenosine methylated mRNAs. Elife 6, (2017).

56. Dember, L. M., Kim, N. D., Liu, K.-Q. & Anderson, P. Individual RNA Recognition Motifs of TIA-1 and TIAR Have Different RNA Binding Specificities. Journal of Biological Chemistry 271, 2783–2788 (1996).

57. Frankish, A. et al. GENCODE 2021. Nucleic Acids Res. 49, D916–D923 (2021).

58. Patro, R., Duggal, G., Love, M. I., Irizarry, R. A. & Kingsford, C. Salmon provides fast and bias-aware quantification of transcript expression. Nat. Methods 14, 417–419 (2017).

59. Soneson, C., Love, M. I. & Robinson, M. D. Differential analyses for RNA-seq: transcript-level estimates improve gene-level inferences. F1000Res. 4, 1521 (2015).

60. Love, M. I., Huber, W. & Anders, S. Moderated estimation of fold change and dispersion for RNA-seq data with DESeq2. Genome Biol. 15, 550 (2014).

61. Dobin, A. & Gingeras, T. R. Mapping RNA-seq Reads with STAR. Curr. Protoc. Bioinformatics 51, (2015).

62. Raudvere, U. et al. g:Profiler: a web server for functional enrichment analysis and conversions of gene lists (2019 update). Nucleic Acids Res. 47, W191–W198 (2019).

63. Dominguez, D. et al. Sequence, Structure, and Context Preferences of Human RNA Binding Proteins. Mol. Cell 70, 854–867.e9 (2018).

64. Luo, X. et al. RMVar: an updated database of functional variants involved in RNA modifications. Nucleic Acids Res. 49, D1405–D1412 (2021).

65. Chen, T. & Guestrin, C. XGBoost. in Proceedings of the 22nd ACM SIGKDD International Conference on Knowledge Discovery and Data Mining 785–794 (ACM, 2016). doi:10.1145/2939672.2939785.

66. Hughes, C. S. et al. Single-pot, solid-phase-enhanced sample preparation for proteomics experiments. Nat. Protoc. 14, 68–85 (2019).

67. Hughes, C. S. et al. Ultrasensitive proteome analysis using paramagnetic bead technology. Mol. Syst. Biol. 10, 757 (2014).

68. Franken, H. et al. Thermal proteome profiling for unbiased identification of direct and indirect drug targets using multiplexed quantitative mass spectrometry. Nat. Protoc. 10, 1567–1593 (2015).

69. Zhang, X. et al. Proteome-wide identification of ubiquitin interactions using UbIA-MS. Nat. Protoc. 13, 530–550 (2018).

70. Gruber, A. R., Lorenz, R., Bernhart, S. H., Neubock, R. & Hofacker, I. L. The Vienna RNA Websuite. Nucleic Acids Res. 36, W70–W74 (2008).

